# Robust lesion network mapping reveals symptom-specific circuits

**DOI:** 10.64898/2026.05.18.725812

**Authors:** Weigang Cui, Jiayi Zhu, Yang Long, Feng Wang, Yilin Yin, Xiongfei Wang, Xiaoxuan Fu, Boyang Zhang, Yuezhu Lu, Wei Zhang, Jianting Huang, Liang Li, Guoming Luan, Yuanxiang Lin, Evan M. Gordon, Danhong Wang, Andrew Zalesky, Nico U.F. Dosenbach, Jianxun Ren, Hesheng Liu

## Abstract

Lesion network mapping (LNM) is a powerful tool for linking distributed brain lesions to symptom-relevant networks, but similar LNM maps across unrelated symptoms have raised concerns about its statistical validity and specificity. Here we introduce robust LNM (rLNM), a statistically principled framework that combines two complementary controls: an anatomy-preserving spatial null model to account for connectome-derived hub bias shared across symptoms, and hypothesis-driven contrasts to resolve selective specificity among related symptoms. rLNM effectively controls both false positives and negatives. Across 10 lesion datasets (384 lesions) and two task-fMRI meta-analyses (339 experiments), rLNM recovers distinct symptom-specific networks (mean *r* = 0.044), which show significantly reduced similarity to the connectome hub pattern (*r* = 0.12 vs. 0.33 in conventional LNM). In 56 patients with neurosurgical lesions, rLNM predicts postoperative language decline at the single-patient level. rLNM thus resolves these concerns and lays a unified statistical foundation for network-symptom inference.

## Introduction

Lesion network mapping (LNM) aims to identify the neural substrates underlying specific neuropsychiatric symptoms by associating anatomically heterogeneous lesions with canonical functional networks^1–8^. In depression, for example, LNM was able to break down the symptoms of a single disorder into two separate networks, one linked to dysphoria and another to anxiosomatic symptoms^9, 10^. LNM has been extended from focal lesions to task activations^4^, atrophy patterns^5^, and brain stimulation sites^7^. As the field continues to mature, recent methodological investigations have raised important questions about the interpretation and specificity of LNM outputs. Some studies have noted that network patterns reported across diverse disorders are similar to each other and converge on hub regions of the normative connectome^11, 12^, while others have demonstrated the existence of symptom-specific lesion networks despite this partial entanglement^13–19^. Together, these observations highlight the need for a rigorous statistical framework that can disentangle genuine, symptom-specific network effects from connectome-derived hub bias, a capacity that conventional LNM (cLNM) does not yet fully provide.

In most implementations, cLNM includes some form of sensitivity and specificity testing^13^. Sensitivity testing identifies voxels connected to a high percentage of lesions associated with a symptom, with a fixed overlap threshold (e.g., ≥75%^20, 21^) applied uniformly across the brain. Since normative connectomes are intrinsically heterogeneous^11^, densely connected hubs, such as the insula, anterior cingulate cortex and frontopolar cortex^22–24^, accumulate lesion-connectivity overlap regardless of the symptom studied. This hub pattern may thus be largely shared across lesion networks, and symptom-specific network effects may be embedded within it. A uniform threshold cannot separate the two, potentially flagging hubs as false positives while missing weaker symptom-specific effects elsewhere. It is also possible that certain symptom or lesion sets do not exhibit a consistent network-level correspondence, such that their spatial distribution is not systematically explained by the connectome. Distinguishing this null scenario from a weak or nonspecific network correspondence is difficult using a uniform threshold alone and requires formal statistical inference. A similar challenge arose in human genetics, where the dominant axes of genotypic variation reflect shared population structure rather than disease, and disease-associated variants became reliably detectable only once statistical methods accounted for this shared background ^25, 26^. LNM likewise calls for a region-specific inference strategy that separates symptom-specific effects from the connectome-derived hub bias, as well as the ability to identify lesion sets for which the connectome provides no evidence of a consistent network-level correspondence. Spatial null models have emerged as a principled solution, evaluating observed convergence against an appropriately constrained null distribution^16, 27^.

Conventional specificity testing in cLNM contrasts the symptom of interest (SOI) against a large, heterogeneous lesion database (e.g., 1,100 lesions spanning 26 symptoms such as aphasia, depression and central pain)^28^. Like task-vs-rest contrasts in fMRI, this foundational approach identifies networks above a diffuse background but does not yet resolve SOI-selective regions from those shared with related symptoms. This distinction is nontrivial, since many symptoms map onto composite neural architectures where shared and SOI-specific contributions are intermixed. For example, motor and visual anosognosia both include unawareness of neurological deficits, but differ in the affected modality (motor vs. visual)^29^. Their lesion networks likely comprise both a shared anosognosia-general component and modality-selective components. This situation parallels a familiar transition in cognitive neuroimaging^30–32^, where task-vs-rest contrasts, though foundational, were later complemented by task-vs-task comparisons (e.g., semantic vs. non-semantic processing^33^) to isolate specific cognitive functions. A similar transition in LNM, from broad-against-unrelated to selective-against-related conditions, thus represents a logical extension for the field.

Here, we developed a robust LNM (rLNM) framework that unifies statistically principled sensitivity and selective specificity testing under a null hypothesis testing framework. Rather than relying on fixed overlap thresholds, rLNM uses an anatomy-preserving spatial null model to test whether a lesion network exists, and further provides explicit statistical control over connectome-derived bias in sensitivity testing. Moreover, rLNM introduces hypothesis-driven selective specificity analyses to isolate symptom-selective lesion networks between mechanistically related conditions. The rLNM framework is applicable to both functional and structural connectomes. We systematically evaluated rLNM using simulated lesions, neuropsychiatric lesions spanning ten symptoms (384 lesions), and task activations from two cognitive domains (339 experiments), as well as a multi-center neurosurgical cohort (56 interventional lesions) with pre- and post-operative language assessments for individual-level causal validation.

## Results

### A unified framework for robust lesion network mapping

First, rLNM implements an anatomy-preserving spatial null model for sensitivity testing (Fig. 1a; see Methods and Supplementary Notes 1-2), similar to the concept proposed by Zalesky and Cash^16^. If a symptom is genuinely underpinned by a specific network, observed lesion-connectivity overlap should exceed what is expected from randomly placed lesions. To construct the null model, each lesion is randomly repositioned across the brain while preserving lesion size, shape, and inter-lesion overlap distribution via simulated annealing^34^. Repeating this procedure 10,000 times generates synthetic lesion network maps, which define a voxel-wise null distribution of lesion-network convergence. The observed convergence is compared voxel-wise with this distribution to obtain an empirical *P* map, with multiple comparisons controlled using the Benjamini–Hochberg false-discovery rate (FDR) correction. Voxels surviving FDR correction (*P*_FDR_ < 0.005; correction across 9,473 whole-brain voxels at 8-mm isotropic resolution) define the null-model-controlled sensitivity map; if no voxel survives, no significant lesion network exists for the lesion set.

**Fig. 1.**
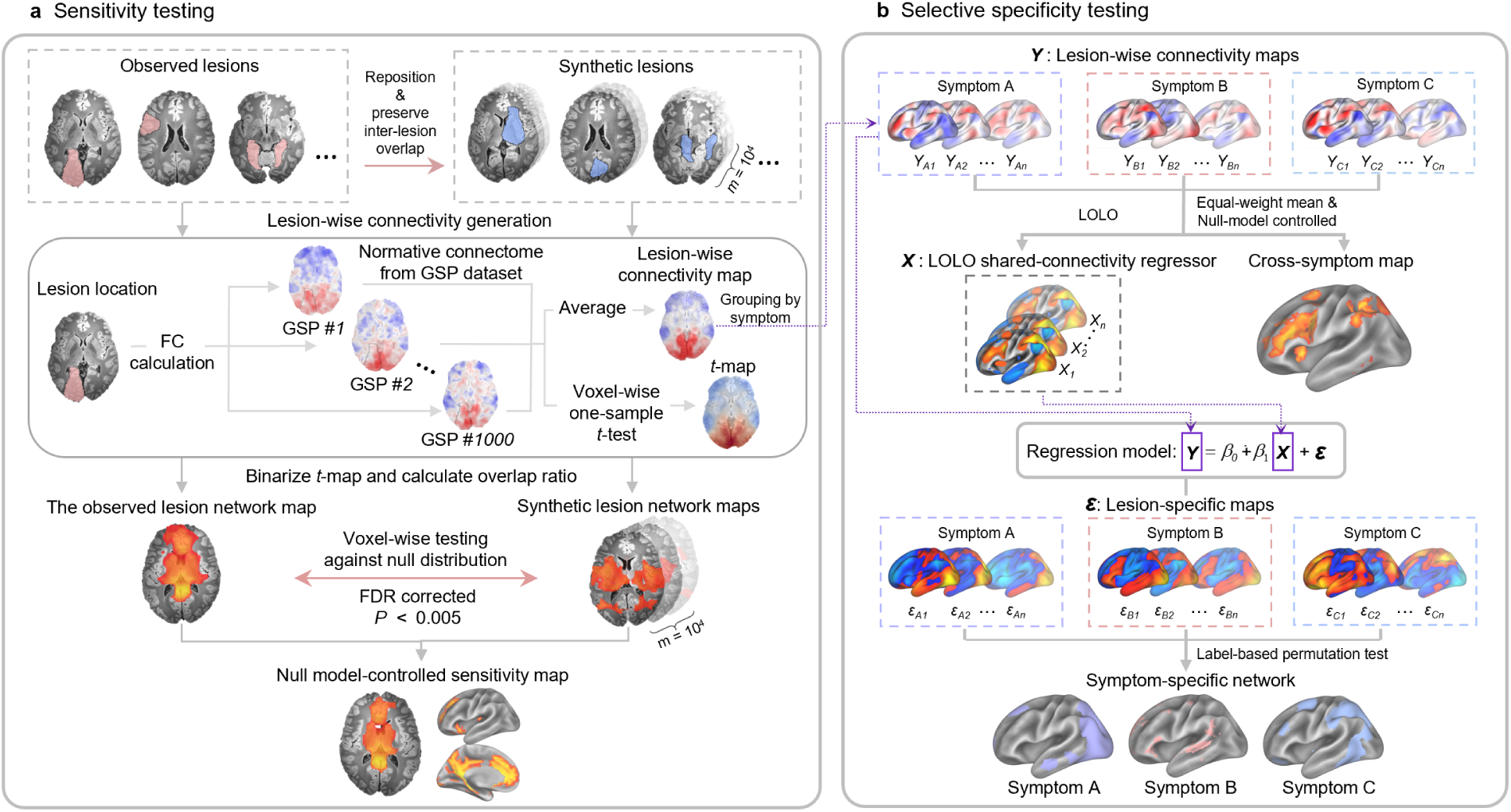
Overview of the robust lesion network mapping framework. a, Sensitivity testing. Observed lesion locations are first used to generate lesion-wise connectivity maps based on normative connectomes derived from 1,000 healthy participants in the GSP dataset. These maps are thresholded and binarized, and their voxel-wise overlap proportion defines the observed lesion network map. Statistical significance is assessed using a spatial null model in which lesions are randomly repositioned while preserving their size, shape and spatial-overlap distribution. Each synthetic lesion set is processed through the same connectivity-mapping pipeline, and 10,000 repetitions generate a voxel-wise null distribution of lesion-network convergence. The observed map is compared with this distribution to identify regions showing greater cross-lesion convergence than expected by chance (*P* < 0.005, false discovery rate (FDR) corrected). **b,** Selective specificity testing. Lesion-wise connectivity maps are first averaged within each symptom. The resulting symptom-level connectivity maps are averaged with equal weight, and then masked by the intersection of all the symptoms’ binary significance masks obtained from null-model sensitivity testing, yielding a cross-symptom map capturing shared connectivity. For symptom-specific analysis, a leave-one-lesion-out (LOLO) shared-connectivity regressor (***X***) is constructed for each lesion. The held-out lesion is excluded when recalculating its symptom-level mean map, and the unthresholded symptom-level maps are then averaged with equal weight across symptoms to obtain the LOLO shared-connectivity regressor. This regressor (***X***) is regressed out of the held-out lesion’s connectivity map (***Y***) to generate a lesion-specific residual map (***ε***). The residual maps are then grouped by symptom, and voxel-wise differences between the symptom of interest and all other symptoms are evaluated by label-based permutation testing. Voxels showing significantly greater residual values for the symptom of interest define the final symptom-specific map.

Second, to isolate networks selective for specific symptoms, rLNM implements selective specificity testing through hypothesis-driven comparisons between related symptoms (Fig. 1b; see Methods). The analysis yields a cross-symptom map capturing shared connectivity and a symptom-specific map for each symptom being compared. To derive the cross-symptom map, lesion-wise connectivity maps are first averaged within each symptom. The resulting symptom-level connectivity maps are then averaged across symptoms and masked by the intersection of all the symptoms’ binary significance masks obtained from null-model sensitivity tests, yielding a cross-symptom map that captures shared connectivity within regions significant for all symptoms. If this intersection is empty, no significant cross-symptom region is identified. Next, symptom-specific effects are isolated using leave-one-lesion-out (LOLO) regression to remove the shared-connectivity component, followed by voxel-wise label-permutation testing. Specifically, each held-out lesion is excluded when recalculating the mean connectivity map for its symptom. The resulting unthresholded symptom-level maps are averaged with equal weight to construct a LOLO shared-connectivity regressor. This regressor serves as the predictor, and the held-out lesion’s unthresholded connectivity map serves as the outcome in a regression across voxels, yielding a lesion-specific residual map. After this procedure is repeated for every lesion, the residual maps are grouped by symptom. At each voxel, residual values for the SOI are contrasted with those for the other symptoms using Welch’s t-tests. Statistical significance was assessed by permuting symptom labels across lesions 10,000 times while preserving the number of lesions assigned to each group. Empirical *P* values were corrected across voxels using the FDR procedure. Voxels with *P*_FDR_ < 0.05 were retained to define the symptom-specific map. Notably, beyond LNM itself, this rLNM framework is applicable to its derivatives, such as activation network mapping^4^ and atrophy network mapping^5, 6^.

### rLNM effectively controls for false positives and false negatives

To test whether rLNM or cLNM better controls connectome hub-driven false positives and threshold-dependent false negatives^11^, we simulated a functional connectivity (FC) matrix consisting of seven large-scale cortical networks with stronger within-network than between-network connectivity, and designated one network as a global hub connected to all others (Supplementary Fig. 1; see Methods). Networks were selected in turn as the target network for detection, and simulated lesions were generated by using lesion-proportion simulation (Fig. 2a; see Methods). Specifically, in each simulated lesion set, a proportion of lesions (*α*) was placed entirely within the target network, while the remaining lesions were randomly distributed across the brain. Varying *α* from 0 to 1, we compared rLNM with cLNM (fixed overlap threshold = 0.75 as in previous studies^20, 21^). Under the complete-null condition (*α* = 0), the estimated family-wise error rate (FWER), defined as the proportion of trials containing at least one false-positive voxel, was 0 for rLNM and 1 for cLNM across 1,000 trials, indicating that rLNM enables rigorous inference on whether a lesion network truly exists. For non-null conditions (*α* > 0), the false-negative rate was the proportion of target-network voxels not identified as significant, with statistical power defined as one minus the false-negative rate. The false-positive rate (FPR) was defined as the proportion of voxels outside the ground-truth network that were incorrectly identified as significant. Both methods could recover the entire target network (statistical power = 1) once *α* reached the 75% overlap threshold. Below this, rLNM consistently outperformed cLNM (Wilcoxon signed-rank test, *P* = 2.44 × 10^-4^), first reaching power = 1 at *α =* 0.65 (Fig. 2b). For false positives, cLNM consistently misidentified the hub network as symptom-relevant (Fig. 2c, FPR of 0.13 ± 0.0058), whereas rLNM’s FPR (0.0008 ± 0.0013) consistently remained lower than that of cLNM across *α* values (*P* = 4.77×10^-7^). Lowering cLNM’s overlap threshold increased statistical power but proportionally inflated FPR, which remained consistently higher than rLNM’s (Supplementary Fig. 2a; all *P* < 10^-3^).

**Fig. 2.**
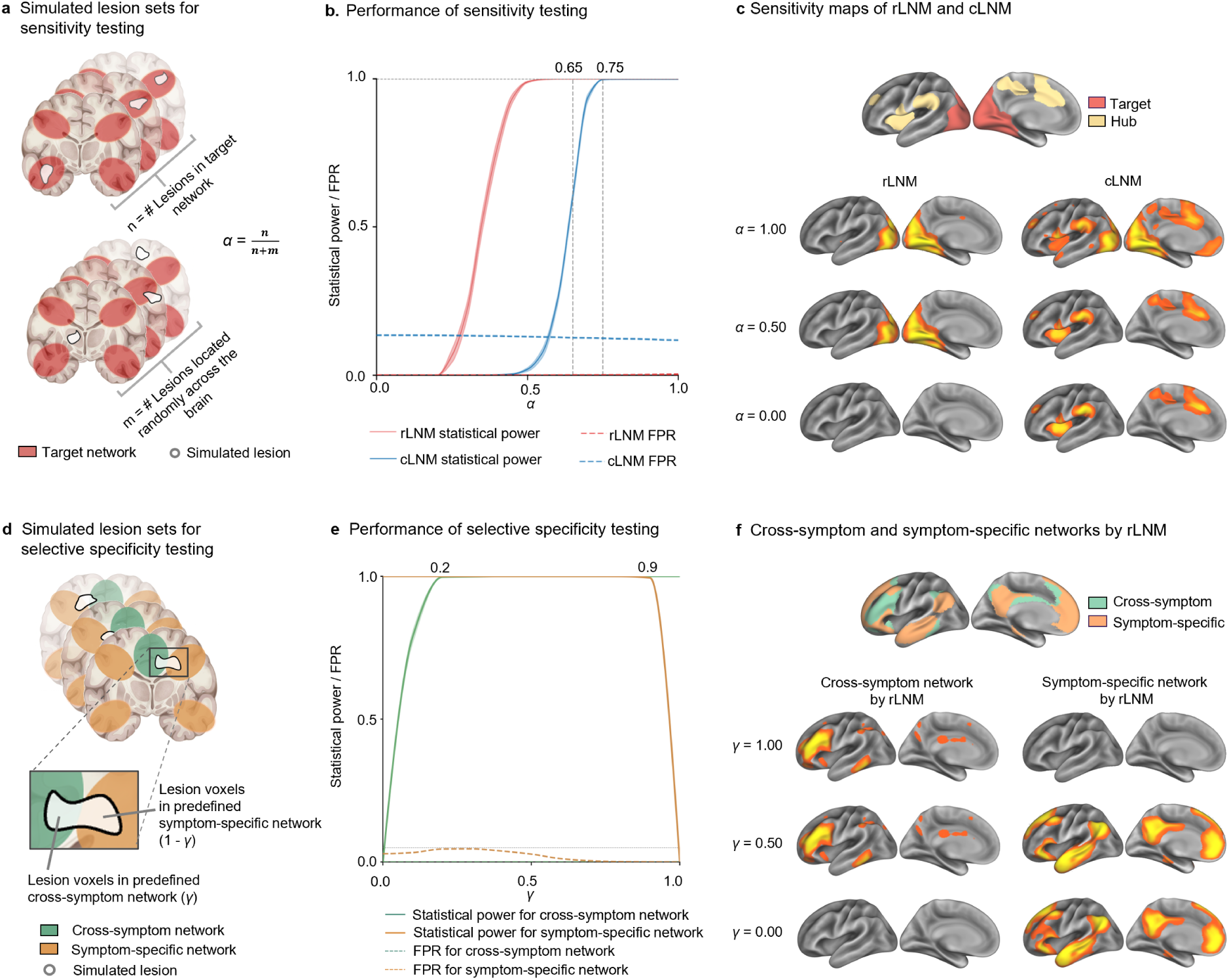
Simulation-based validation of false-positive and false-negative control in rLNM. **a**, Generation of simulated lesion sets in the lesion-proportion simulation. Each lesion set comprised *n* lesions located entirely within the target network and *m* lesions randomly positioned across the whole brain, yielding a target-lesion proportion of *α* = *n*/(*n* + *m*). **b**, Statistical power (solid lines) and false-positive rate (FPR; dotted lines) of rLNM (red) and cLNM (blue) across *α* values ranging from 0 to 1. Statistical power reaches 1 at *α* = 0.65 for rLNM and *α* = 0.75 for cLNM. Across α values, the FPR of rLNM (0.0008 ± 0.0013) is consistently lower than that of cLNM (0.13 ± 0.0058; Wilcoxon signed-rank test, *P* = 4.77×10^-7^). **c**, Sensitivity maps obtained using rLNM and cLNM at representative *α* values. The top row shows the predefined hub and target networks. Subsequent rows show sensitivity maps obtained using rLNM (left) and cLNM (right) at *α* = 1.0, 0.5, and 0. Hub regions are apparent in all maps derived from cLNM but are well controlled by rLNM. **d**, Generation of simulated lesion sets in the shared-specific simulation. In each simulated lesion (gray outline), a proportion *γ* of voxels was placed within the predefined cross-symptom network, and the remaining proportion, 1-*γ*, within the symptom-specific network. **e**, Statistical power (solid lines) and FPR (dotted lines) of the cross-symptom network (green) and symptom-specific network (orange) across *γ* values ranging from 0 to 1. The horizontal dashed line indicates a reference FPR of 0.05. Statistical power reaches 1 for *γ ≥* 0.20 in the cross-symptom network and *γ* ≤ 0.90 in the symptom-specific network. The FPR remains well below 0.05 across *γ* values for both the cross-symptom network (0.00007 ± 0.00022) and the symptom-specific network (0.021 ± 0.0184; Wilcoxon signed-rank test, both *P* < 10^-3^). **f**, Selective specificity maps obtained at representative *γ* values. The top row shows the predefined cross-symptom and symptom-specific networks. Subsequent rows show cross-symptom maps and symptom-specific maps obtained using rLNM selective specificity testing at *γ* = 1, 0.5, and 0. Values are mean ± s.d. across simulation trials.

We next used voxel-proportion simulation (Extended Data Fig. 1a; see Methods), in which a proportion *β* of voxels in each simulated lesion fell within the target network and the remaining voxels were distributed elsewhere. Under the complete-null condition (*β* = 0), the estimated FWER was 0 for rLNM and 1 for cLNM across 1,000 trials. rLNM reached statistical power = 1 at *β* = 0.25 and consistently outperformed cLNM below *β* = 0.35 (Wilcoxon signed-rank test, *P* = 0.03; Extended Data Fig. 1b). By mitigating connectome-derived hub bias (Extended Data Fig. 1c), rLNM maintained a significantly lower FPR (0.0023 ± 0.0037) than cLNM (0.14 ± 0.0008; *P* = 4.77×10^-7^) across *β* values. Sweeping cLNM’s overlap threshold from 45% to 85% revealed a consistent power-FPR trade-off, with cLNM’s FPR remaining significantly higher than rLNM’s at every threshold (Supplementary Fig. 2b; all *P* < 10^-3^).

For selective specificity testing, we constructed simulations that combined cross-symptom and symptom-specific networks (Fig. 2d; see Methods). In each simulated lesion, a proportion of voxels (*γ*) was placed within the predefined cross-symptom network and the remaining voxels within the symptom-specific network. At *γ* = 1, when no symptom-specific component was present, the FWER for symptom-specific testing was 0.003. Across *γ* values ranging from 0 to 1, FPRs for both the cross-symptom network (0.00007 ± 0.00022) and symptom-specific network (0.021 ± 0.018) remained well below 0.05 (Wilcoxon signed-rank test, both *P* < 10^-3^; Fig. 2e). Statistical power reached 1 for the cross-symptom network at *γ* ≥ 0.2 and for the symptom-specific network at *γ* ≤ 0.9 (Fig. 2e, f), confirming effective control of both error types.

### rLNM identifies biologically meaningful, symptom-specific lesion networks

To evaluate rLNM’s performance on empirical lesions, we applied both rLNM and cLNM to ten publicly available lesion datasets spanning diverse neuropsychiatric symptoms (Supplementary Table 1). No voxels survived rLNM null-model inference in the mixed-lesion dataset (*n* = 100; randomly sampled across ten clinical symptoms; see Methods), the migraine dataset (*n* = 11), or the stuttering dataset (*n* = 20; Fig. 3a, left; Extended Data Fig. 2; see significant voxel counts in Supplementary Table 2), indicating no genuine underlying lesion network in these lesion sets, or that the lesion sets are too small in sample size or too noisy to enable identification of a lesion network. By contrast, cLNM showed substantial lesion-network convergence in hub regions such as the insula and dACC (Fig. 3a, right; Extended Data Fig. 3). These maps closely correspond to the summation vector of the normative connectome (sum of *C*)^11^ (*r* = 0.461, 0.560 and 0.529, respectively).

**Fig. 3.**
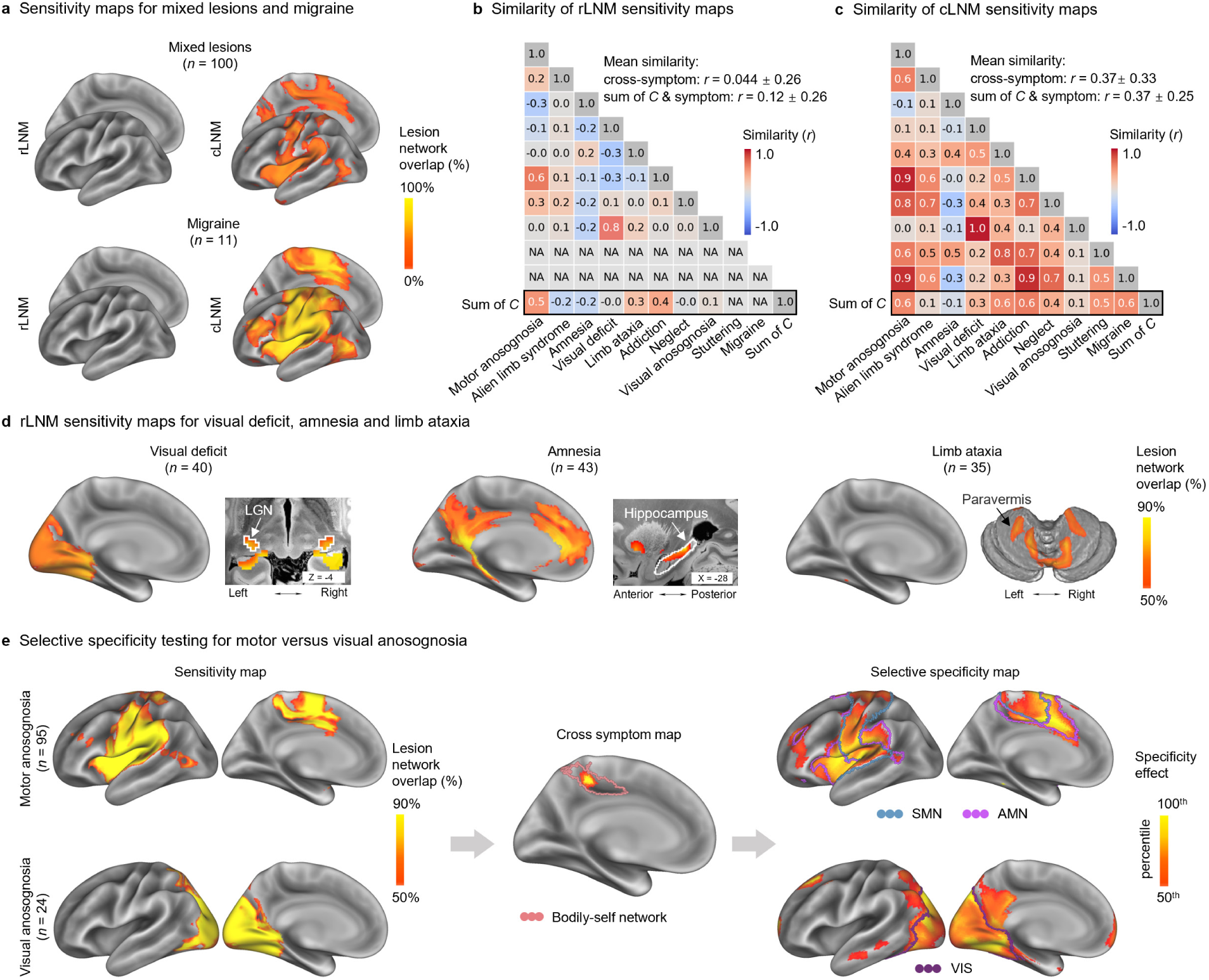
rLNM achieves symptom-specific network mapping across neuropsychiatric symptoms. **a**, rLNM and cLNM sensitivity maps derived from mixed lesions (*n* = 100) and migraine (*n* = 11). No region survived rLNM null-model inference in either dataset, whereas cLNM showed convergence in dACC and insula (see whole-cortex maps of rLNM and cLNM in Extended Data Fig. 2, 3). **b**, Similarity of rLNM sensitivity maps. No significant lesion network was identified for migraine or stuttering under null-model inference (*P*_FDR_ < 0.005). Across the remaining symptoms, rLNM maps were dissimilar across symptoms (cross-symptom similarity, *r* = 0.044 ± 0.26), and showed little correspondence with the sum of *C* (*r* = 0.12 ± 0.26). **c**, Similarity of cLNM sensitivity maps. cLNM maps showed moderate similarity across symptoms (*r* = 0.37 ± 0.33), and resembled the summed connectivity profile of the normative connectome (sum of *C*; *r* = 0.37 ± 0.25). **d**, rLNM sensitivity maps across three representative symptoms (see corresponding cLNM maps in Extended Data Fig. 3). The map for visual deficit (*n* = 40) highlights visual cortex (spin test, 1,000 permutations, one-sided *P*_spin_ < 10^-3^; *P*_spin_ > 0.05 for all other networks) and lateral geniculate nucleus (spatial permutation test, 1,000 permutations, one-sided *P*_perm_= 0.008). The map for amnesia (*n* = 43) identifies the anterior/posterior cingulate cortex (all *P*_spin_ < 10^-3^) and hippocampus (*P*_perm_ = 0.038). The map for limb ataxia (*n* = 35) highlights the paravermis (*P*_perm_ = 0.039). **e**, Dissociation of cross-symptom and symptom-specific effects derived from motor anosognosia (*n* = 95; top) and visual anosognosia (*n* = 24; bottom). rLNM sensitivity maps are shown on the left. Their cross-symptom map (middle) overlaps with the bodily-self network (pink boundary, *P*_spin_ = 0.007). Specificity maps (right) isolate symptom-specific effects after accounting for the shared component: motor anosognosia preferentially involves the somatomotor network (SMN; blue boundary) and action-mode network (AMN, purple boundary; all *P*_spin_ < 10^-3^), whereas visual anosognosia preferentially involves the visual network (VIS, purple boundary; *P*_spin_ < 10^-3^).

We then quantified pairwise spatial similarity between symptom sensitivity maps and compared all of them to the sum of *C* (Extended Data Fig. 4a). Under rLNM, migraine and stuttering yielded empty sensitivity maps and were therefore excluded from the spatial-correlation analysis. Across the remaining eight lesion datasets, rLNM maps showed low between-symptom similarity (*r* = 0.044 ± 0.26) and limited similarity to the sum-of-*C* pattern (*r* = 0.12 ± 0.26; Fig. 3b). Both were significantly lower than the corresponding values for cLNM (between-symptom similarity: *r* = 0.37 ± 0.33; Wilcoxon signed-rank test, *P* = 3.20 × 10^-7^; Fig. 3c and Extended Data Fig. 4b; similarity to sum-of-*C* pattern: *r* = 0.37 ± 0.25; *P* = 7.81 × 10^-3^; Extended Data Fig. 4c). Importantly, biologically interpretable symptom-specific networks were preserved in rLNM maps. For example, the sensitivity map of visual deficit (*n* = 40) predominantly localized to the visual cortex (Fig. 3d; spin test, 1,000 permutations, one-sided *P*_spin_ < 10^-3^; *P*_spin_ > 0.05 for all other networks) and lateral geniculate nucleus (spatial permutation test, 1,000 permutations, one-sided *P*_perm_ = 0.008), a principal thalamic relay of the retinocortical visual pathway^35^. The amnesia (*n* = 43) map highlighted the anterior/posterior cingulate cortex (all *P*_spin_ < 10^-3^) and hippocampus (*P*_perm_ = 0.038), which are closely associated with memory function^36^. In limb ataxia (*n* = 35), significant effects localized predominantly to the cerebellar paravermis (one-sided spatial permutation test, *P*_perm_ = 0.039) rather than cerebral cortex, corresponding to sensorimotor cerebellar territories recruited during limb movements^37^. Other lesion sets likewise yielded anatomically circumscribed patterns rather than broadly distributed whole-brain networks, including marked right lateralization in neglect^38^ and convergence on the pars marginalis of the cingulate sulcus^39, 40^ in alien limb syndrome (Extended Data Fig. 2).

To demonstrate how selective specificity distinguishes shared from symptom-specific components, we examined motor (*n* = 95) and visual anosognosia (*n* = 24)^29^, two symptoms sharing an anosognosia-general impairment but in different modalities. Based on the sensitivity maps for each symptom (Fig. 3e, left), we first derived a cross-symptom map capturing their shared network pattern (Fig. 3e, middle). This map converged on the pars marginalis of the cingulate sulcus, overlapping with the recently defined bodily-self network^39^ (one-sided *P*_spin_ = 0.007; *P*spin > 0.05 for all other networks; Supplementary Fig. 3a), which integrates body schema information with ongoing action to maintain an updated sense of the bodily self, as further supported by electrical stimulation^40^. We next derived specificity maps for each symptom (Fig. 3e, right), which predominantly involved the corresponding action (including action-mode and somatomotor networks) and visual systems (all *P*_spin_ < 10^-3^; *P*_spin_ > 0.05 for all other networks; Supplementary Fig. 3b, c), consistent with their distinct functional symptoms.

### rLNM identifies canonical and task-specific functional networks from task activations

Because the ground-truth functional substrates underlying lesion-induced symptoms are inherently unknown, we further benchmarked our robust approach using activation network mapping^4^, a derivative of LNM in which task activations serve as foci. Task activations converge on canonical functional networks in task fMRI, providing an independent reference for assessing whether rLNM recovers the corresponding networks. Task-based fMRI activation coordinates were sourced from the BrainMap database (see Methods).

In sensitivity testing, rLNM isolated the language network (spin test, 1,000 permutations, one-sided *P*_spin_ < 10^-3^; Fig. 4a, left; Extended Data Fig. 5a), with no significant involvement of any other network (all *P*_spin_ > 0.05), demonstrating that rLNM achieved task-specific network mapping. By contrast, cLNM yielded a widespread pattern for language tasks (*n* = 155 experiments; Fig. 4a, right; Extended Data Fig. 5b), including the language, auditory and action-mode networks (*P*_spin_ < 10^-3^ for both the language and auditory networks and *P*spin = 0.015 for the action-mode network).

**Fig. 4.**
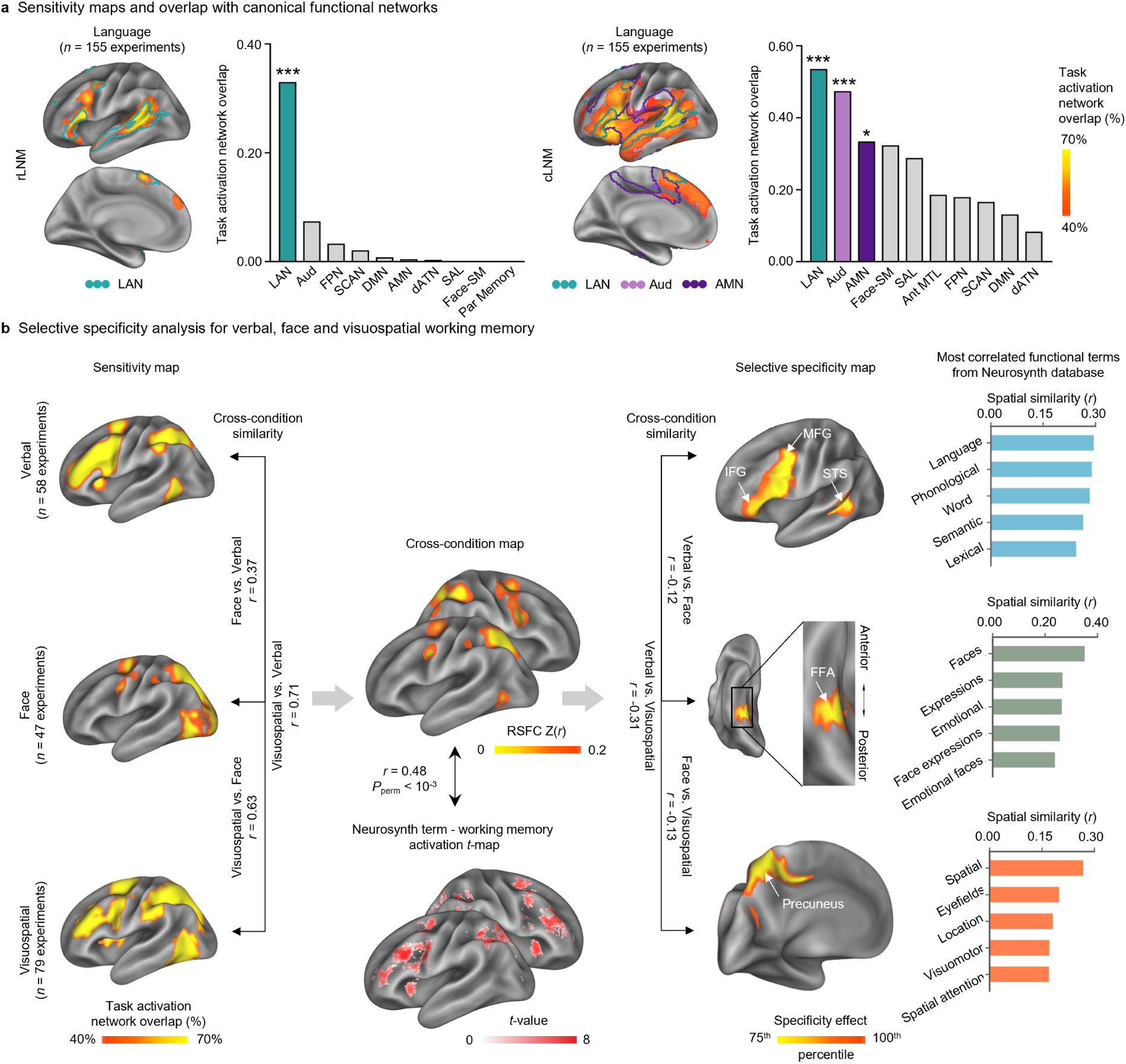
rLNM achieves condition-specific network mapping across cognitive domains. a, Cortical visualizations of rLNM and cLNM sensitivity maps for language tasks (*n* = 155 experiments). The rLNM map was specific to the language network (spin test, 1,000 permutations, one-sided *P*_spin_ < 10^-3^; all *P*_spin_ > 0.05 for other networks). By contrast, cLNM highlighted the language (*P*_spin_ < 10^-3^), auditory (*P*_spin_ < 10^-3^) and action-mode (*P*_spin_ = 0.015) networks. **b**, Dissociation of cross-condition and condition-specific patterns across working-memory conditions. Based on the sensitivity maps (left) from verbal (*n* = 58 experiments), face (*n* = 47 experiments) and visuospatial (*n* = 79 experiments) working memory (WM), the resulting cross-condition map (middle) shows significant spatial overlap with the Neurosynth working-memory activation map (*r* = 0.48, *P*_perm_ < 10^-3^). Selective-specificity maps (right) reveal distinct patterns for each condition. The verbal WM map highlights regions of inferior frontal gyrus (IFG; *P*_spin_ < 10^-3^), superior temporal sulcus (STS; *P*_spin_ = 0.036), and posterior middle frontal gyrus (pMFG; *P*_spin_ = 0.046). The face and visuospatial WM maps highlight fusiform face area (FFA, *P*_spin_ = 0.019), and precuneus (*P*_spin_ = 0.038), respectively. Bar plots show the Neurosynth terms most strongly correlated with each selective-specificity map. *** *P* < 10^-3^. * *P* < 0.05.

For selective specificity analyses, we collected activation coordinates from three types of working memory (WM) tasks: verbal (*n* = 58 experiments), face (*n* = 47 experiments) and visuospatial (*n* = 79 experiments; see sensitivity maps in Fig. 4b, left). The cross-condition map generated by aggregating these three tasks strongly overlapped with the ‘working memory’ map derived from the Neurosynth database (*r* = 0.48, *P*_perm_ < 10^-3^; Fig. 4b, middle). We then derived condition-specific maps by regressing out the cross-condition component, substantially reducing the high inter-map similarity observed prior to regression (pairwise *r* = 0.37, 0.63, and 0.71), resulting in maps with minimal spatial overlap (pairwise *r* = -0.12, -0.13, and -0.31; Fig. 4b, right). Verbal WM effects were predominantly localized to the inferior frontal gyrus (IFG, *P*_spin_ < 10^-3^), superior temporal sulcus (STS, *P*_spin_ = 0.036) and posterior middle frontal gyrus (pMFG; *P*_spin_ = 0.046), all within the language network. Face WM effects were enriched in the fusiform face area (FFA; *P*_spin_ = 0.019), while visuospatial WM effects were enriched in the precuneus (*P*_spin_ = 0.038). This dissociation is consistent with the involvement of the distributed frontotemporal language network in language processing^41^, the FFA in face-selective processing^42^, and the precuneus in spatial representation and manipulation^43^. The most associated Neurosynth terms were ‘language’ (*r* = 0.29), ‘faces’ (*r* = 0.35), and ‘spatial’ (*r* = 0.26; one-sided spatial permutation test, all *P*_perm_ < 10^-3^) for the verbal, face, and visuospatial WM, respectively, indicating that the specificity testing effectively captured task-specific effects.

### rLNM predicts postoperative language decline at the individual level

Epilepsy neurosurgery provides a unique opportunity for causal validation of rLNM, offering anatomically precise resections paired with pre- and post-operative behavioral assessments at the individual level. We thus tested whether an rLNM-derived language network could predict postoperative language decline in individual patients. We recruited a total of 56 patients (24 females; age = 25.7 ± 9.6 years) with drug-resistant epilepsy from two clinical centers (Fig. 5a). All patients underwent structural MRI and Western Aphasia Battery (WAB) assessment within 1 week before surgery and again 1 week postoperatively. Based on postoperative WAB change (see Methods), patients were classified into aphasia (*n* = 29) and non-aphasia (*n* = 27) groups. The groups did not differ significantly in age, sex distribution, or lesion volume (Supplementary Fig. 4). Lesion distributions showed modest spatial similarity between the two groups (*r* = 0.26, CI = [0.25, 0.27], *P*_spin_ = 0.009; Fig. 5a, rightmost column).

**Fig. 5.**
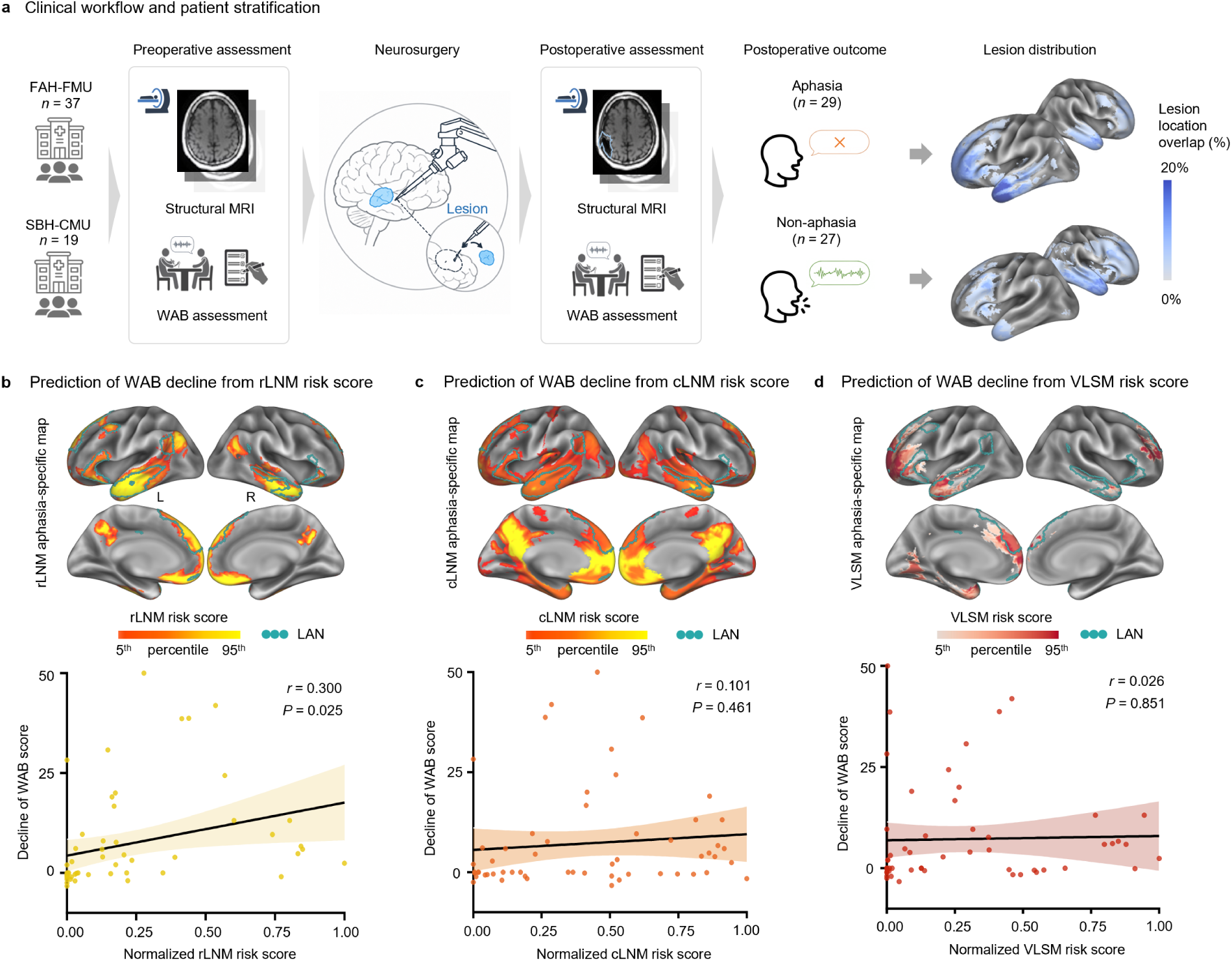
Patient-level interventional validation of rLNM in epilepsy surgery. **a**, Clinical workflow and patient stratification. Patients with epilepsy were recruited from two clinical centers (FAH-FMU, *n* = 37; SBH-CMU, *n* = 19) and underwent structural MRI and Western Aphasia Battery (WAB) assessment before and after epilepsy surgery. Based on postoperative WAB change (see Methods), patients were classified into aphasia (*n* = 29) and non-aphasia (*n* = 27) groups. The lesion distributions of the two groups showed modest spatial similarity (*r* = 0.26, CI = [0.25, 0.27], *P*_spin_ = 0.009). **b**, The rLNM-derived aphasia-specific map aligns well with the language network (LAN; cyan), and rLNM-derived risk scores significantly predicted postoperative WAB change (*r* = 0.300, CI = [0.040, 0.52], *P* = 0.025). **c**, The cLNM-derived aphasia-specific map highlighted the DMN and LAN. cLNM-derived risk scores did not predict postoperative WAB change (*r* = 0.101, CI = [-0.17, 0.35], *P* = 0.461). **d**, The aphasia-specific map derived from voxel-based lesion-symptom mapping (VLSM) predominantly highlights frontal regions and shows limited correspondence with the LAN. The scatter plot shows that the VLSM-derived risk scores, quantifying the extent of each surgical lesion’s involvement in the map, failed to predict postoperative WAB change (*r* = 0.026, *P* = 0.851). Brain maps are displayed from the 5^th^ to the 95^th^ percentile of map values. Shaded regions indicate 95% confidence intervals.

The rLNM-derived aphasia-specific map primarily involved the language network (*P*_spin_ < 10^-3^; Fig. 5b). The extent of each surgical lesion’s involvement in this aphasia-specific network (defined as the risk score; see Methods) significantly predicted the magnitude of postoperative WAB decline (leave-one-subject-out cross-validation [LOSOCV]; *r* = 0.300, CI = [0.04, 0.52], *P* = 0.025). By contrast, the cLNM-derived aphasia-specific map identified a broad lesion network highlighting the DMN (*P*_spin_ < 10^-3^), followed in prominence by the language network (*P*_spin_ = 0.002; Fig. 5c). The corresponding risk scores did not significantly predict postoperative WAB changes (LOSOCV; *r* = 0.101, CI = [−0.17, 0.35], *P* = 0.461).

As an additional control, we tested whether focal lesion anatomy could account for postoperative language decline. Voxel-based lesion-symptom mapping (VLSM; see Methods) contrasting the aphasia and non-aphasia groups predominantly identified frontal regions associated with postoperative decline (Fig. 5d). This pattern showed greatest correspondence with the limbic network (*P*_spin_ = 0.012), rather than the language network (*P*_spin_ = 0.062), and did not significantly predict postoperative WAB changes (LOSOCV; *r* = 0.026, CI = [−0.24, 0.29], *P* = 0.851). Prediction remained non-significant when using the label-permutation-corrected VLSM map (*r* = −0.031, CI = [−0.29, 0.23], *P* = 0.818; Extended Data Fig. 6a). Direct overlap between each resection and the canonical language network likewise did not significantly predict postoperative WAB changes (*r* = 0.045, CI = [−0.22, 0.30], *P* = 0.744; Extended Data Fig. 6b). Together, these results provide patient-level interventional validation of rLNM, showing that an rLNM-derived network can predict the functional consequences of focal brain injury beyond lesion anatomy alone.

### Structural rLNM reveals symptom-specific white-matter pathways

We next investigated whether the rLNM framework could be extended from functional to structural connectivity to identify symptom-specific white-matter pathways (see Methods). We applied structural-connectome-based rLNM to the ten symptoms examined above (Fig. 6a; Fig. 6b, left; Extended Data Fig. 7). In sensitivity testing, rLNM identified the fornix, a principal hippocampal pathway critically involved in episodic memory^44^, as a major pathway associated with amnesia (*n* = 43; one-sided spatial permutation test, 1,000 permutations, *P*_perm_ < 10^-3^). For limb ataxia (*n* = 35), no cerebral white-matter pathway survived significance testing, whereas the predominant tract identified was the superior cerebellar peduncle (SCP, *P*_perm_ < 10^-3^), a major cerebellar efferent pathway involved in coordinated motor control^45^ whose degeneration is associated with greater ataxia severity^46^. For alien limb syndrome (*n* = 48), significant effects localized to the corpus callosum (*P*_perm_ < 10^-3^), consistent with its role in interhemispheric sensorimotor integration and with previous evidence linking callosal disconnection to alien limb phenomena^47, 48^. Across the remaining symptoms, rLNM identified biologically plausible symptom-relevant pathways (Extended Data Fig. 7; Supplementary Table 3), including the right frontal aslant tract (FAT, *P*_perm_ = 0.008) for motor anosognosia^49^, the right superior longitudinal fasciculus II (SLF II, *P*_perm_ = 0.003) for spatial neglect^50^, the left frontal aslant tract (FAT) and arcuate fasciculus (AF, *P*_perm_ = 0.003) for stuttering^51–53^, and the corticostriatal pathway (*P*_perm_ < 10^-3^) for addiction^54, 55^. No white-matter pathway reached significance for migraine. By comparison, cLNM produced substantially broader tract patterns (Extended Data Fig. 8), including the pathways identified by rLNM but also additional tracts with less clear relevance to the corresponding symptoms, such as the corpus callosum for amnesia (*P*_perm_ < 10^-3^) and the optic radiation for stuttering (*P*_perm_ < 10^-3^).

**Fig. 6.**
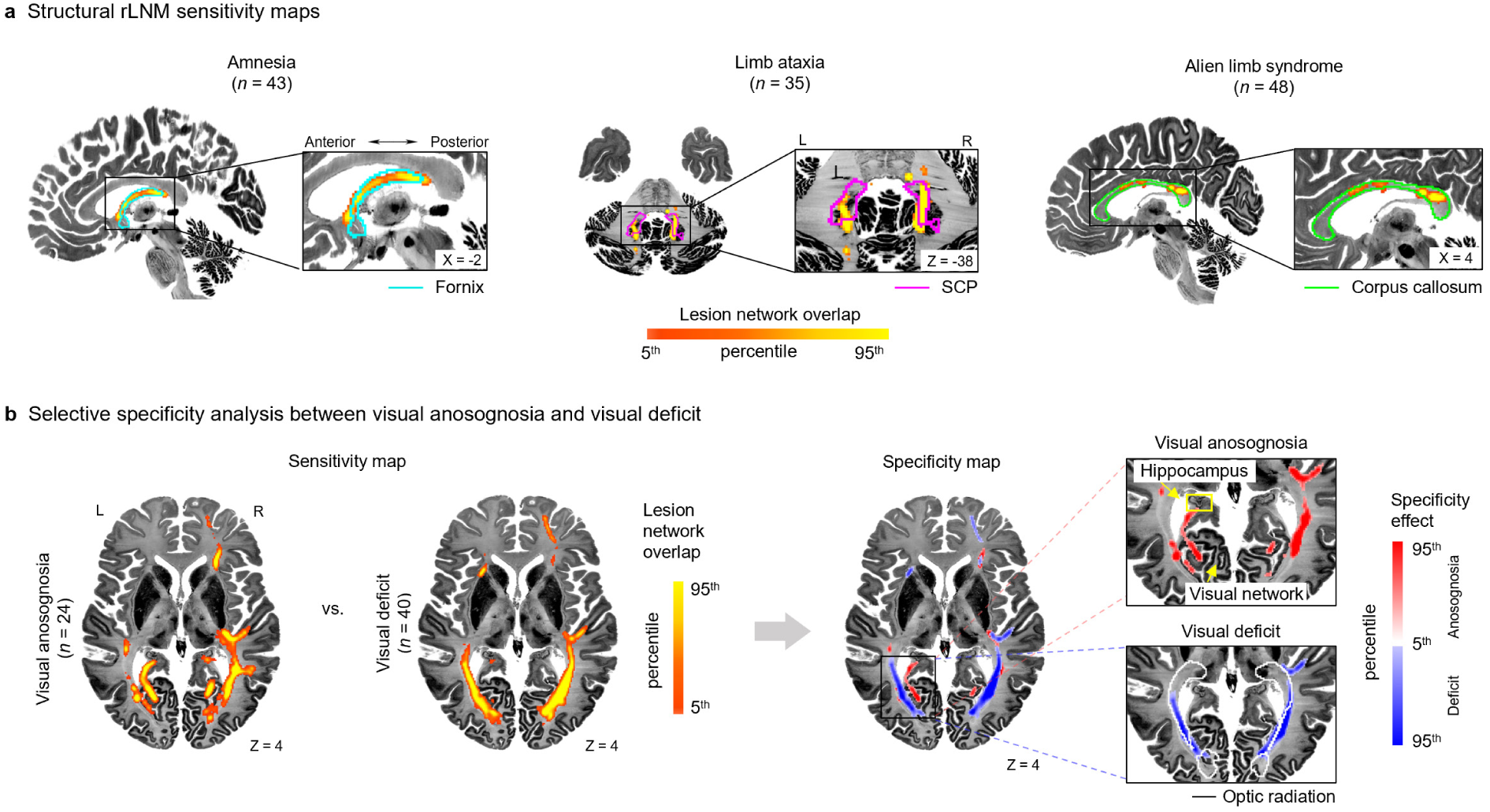
Structural rLNM identifies symptom-relevant and symptom-selective white-matter pathways. a, Structural rLNM sensitivity maps for amnesia (*n* = 43), limb ataxia (*n* = 35) and alien limb syndrome (*n* = 48; see corresponding cLNM maps in Extended Data Fig. 8). Warm-colored overlays indicate lesion structural connectivity overlap, displayed as percentile ranks from the 5^th^ to the 95^th^ percentile. Representative white-matter tracts are outlined for anatomical reference: the fornix (one-sided spatial permutation test, *P*perm < 10^-3^; cyan) for amnesia, the superior cerebellar peduncle (SCP, *P*_perm_ < 10^-3^; magenta) for limb ataxia and the corpus callosum (*P*_perm_ < 10^-3^; green) for alien limb syndrome. **b,** Selective specificity testing comparing visual anosognosia (*n* = 24) and visual deficit (*n* = 40). Sensitivity maps for each symptom are shown on the left, with warmer colors indicating higher overlap percentiles. The specificity map identifies tracts preferentially associated with visual anosognosia (red) or visual deficit (blue), with color intensity indicating the percentile-ranked specificity effect. Zoomed-in views highlight preferential involvement of a visual-hippocampal pathway (*P*_perm_ < 10^-3^) in visual anosognosia and the optic radiation (*P*_perm_ < 10^-3^) in visual deficit.

We next examined selective specificity between visual anosognosia and visual deficit using structural rLNM. In the sensitivity maps, both symptoms involved white-matter pathways associated with the visual system (Fig. 6b, left). Selective specificity analysis, however, revealed distinct pathways for the two symptoms (Fig. 6b, right). Visual deficit preferentially involved the optic radiation (*P*_perm_ < 10^-3^), the major pathway transmitting visual information from the lateral geniculate nucleus (LGN) to the visual cortex, whose disruption is a well-established cause of visual field loss^56^. By contrast, visual anosognosia preferentially involved a white-matter pathway linking the visual system with the hippocampus (*P*_perm_ < 10^-3^). The hippocampus has been implicated in comparing current sensory input with memory-based expectations^57, 58^, so this pathway may be particularly relevant to sensory awareness.

## Discussion

### Spatial null model separates symptom-specific networks from shared connectome background

Recent methodological work^11, 12, 59^ has shown that some LNM maps may be linear combinations of rows of the normative connectome, whose variance is dominated by a few low-dimensional gradients. Maps derived for unrelated conditions can therefore resemble one another and the normative connectome (sum of *C*), an observation our cLNM analyses also reproduce (Fig. 3a, c). However, high similarity between maps does not rule out meaningful differences between them. As illustrated intuitively by Zalesky and Cash^16^ (Fig. 1g in ref. 16), two maps can remain highly correlated even when they differ clearly within a focal region. Likewise, lesions associated with different symptoms sample different parts of the connectome, and the resulting deviations may explain little variance while still carrying symptom-specific information. As in human genetics, where disease-associated variants become detectable only after accounting for shared population structure^25, 26^, the central question is not how much variance such signals explain, but whether they can be reliably separated from the shared background.

rLNM addresses this question with an anatomy-preserving spatial null model. Since null lesion sets are drawn from the same connectome and matched to the observed lesions in number, size, shape and overlap, the shared background is present in both observed and null maps. This replaces a uniform overlap threshold with region-specific inference, in which each region is tested against the overlap expected by chance at that location, so hubs require higher overlap to reach significance. rLNM thereby controlled hub-driven false positives in simulations (Fig. 2) and found no significant voxels in mixed-lesion, migraine, and stuttering datasets whose cLNM maps closely resembled the normative connectome (sum of *C*) (Fig. 3). These null findings illustrate that rLNM requires sufficient evidence of convergence beyond null expectations; in migraine (*n* = 11) and stuttering (*n* = 20), such evidence may be limited by small sample sizes or the absence of a common underlying functional network. Where significant convergence was detected, some resulting maps showed symptom-specific patterns that were more refined than canonical large-scale networks: a right-lateralized network for spatial neglect, a sensorimotor cerebellar network for limb ataxia, and a circumscribed region for alien limb syndrome (Fig. 3, Extended Data Fig. 2). Such lateralized and focal maps are difficult to express with whole-brain, largely symmetric connectome gradients, consistent with the view that lesion-specific sampling of the connectome carries information beyond its dominant low-dimensional gradients^16^.

Specificity testing in cLNM¹² follows a similar logic, contrasting target lesions with a large, heterogeneous cohort that effectively serves as an empirical null. rLNM can be viewed as its *in silico* counterpart, replacing this cohort with the same lesions placed at random. This replacement matters because an empirical cohort comes from different patients and etiologies, most often stroke, and can therefore differ from the target lesions in size, location, and lesion overlap. rLNM avoids these differences because its null lesions are the same lesions placed at random, so only convergence beyond chance remains significant. Practically, the *in silico* reference also removes the need for a large lesion cohort, which is rarely publicly available, making rigorous inference accessible to any rLNM analysis.

### Selective specificity separates shared from symptom-specific brain patterns

Many neurological and psychiatric conditions comprise multiple co-occurring dysfunctional components supported by distinct neural systems^9, 60, 61^. To disentangle these components, we introduce ’selective specificity’ as a new inferential concept in rLNM. Unlike the aforementioned specificity in cLNM, selective specificity establishes a paradigm for well-controlled, hypothesis-driven inference among related conditions. This conceptual framework echoes the late-1990s shift in cognitive neuroimaging from generic task-versus-rest comparisons to carefully matched task-versus-task contrasts^30, 33^, which attempted to separate shared task-related activity from process-specific components. For example, contrasting verbal against spatial working memory reveals a shared frontoparietal control network alongside modality-specific contributions in left frontal/temporal cortex for verbal processes, and right parietal cortex for spatial processes^62^. Likewise, selective specificity enables LNM to decompose a symptom’s network map into shared and symptom-specific components. Under this view, both our rLNM implementation and prior studies^29, 61, 63^ that directly contrasted related conditions can be seen as instances of a broader selective-specificity framework. Beyond hypothesis-driven comparisons, as exemplified above, data-driven exploration is also encouraged to uncover unexpected shared components across seemingly unrelated disorders, although such findings require rigorous external validation and cautious interpretation.

More broadly, selective specificity addresses a general inferential problem in brain mapping: a spatial pattern associated with a condition is not necessarily selective to it. The conventional case-control design in brain mapping, which compares patients against healthy controls, establishes association with the disease but not selectivity relative to other disorders, and may therefore misinterpret shared cross-disorder abnormalities as condition-specific features. For example, large-scale case-control studies of major depressive disorder (MDD) have identified cortical abnormalities in the anterior cingulate and insular cortex^64^. However, transdiagnostic analyses showed that gray-matter reductions in these regions are shared across multiple psychiatric disorders, while more pronounced anterior hippocampal and amygdala reductions are selective to MDD^65^. A parallel issue arises in neuroimaging transcriptomics, where spatial neuroimaging phenotypes are correlated with normative gene-expression patterns to infer their molecular correlates^66^. Because cortical gene expression follows broad macroscale organizational gradients^67^, an observed neuroimaging-transcriptomic association may reflect a mixture of general cortical organization and phenotype-specific molecular variation. The selective-specificity principle and its statistical implementation address this problem by introducing biologically meaningful comparison maps to distinguish shared from phenotype-specific components. This framework is therefore broadly applicable across pattern-based neuroimaging analyses, including transdiagnostic brain mapping^65^, imaging-transcriptomic mapping^68^ and other emerging approaches that seek to infer biological mechanisms from spatial correspondence.

### rLNM extends robust network inference to structural pathways

Although LNM has been implemented predominantly using functional connectivity, structural connectomes have been incorporated both into lesion-based network mapping^69–73^ and stimulation-based fiber mapping^74^, providing tract-level information about symptom-relevant white-matter pathways. This tract-level network mapping, however, introduces a distinct statistical challenge that is less pronounced in functional LNM. Structural LNM is particularly sensitive to lesion size and shape: larger lesions generally intersect more streamlines, and lesion shape can substantially influence which white-matter pathways are affected^70^. Consequently, observed tract convergence does not fully reflect a tract’s genuine involvement in the symptom, but is partly driven by lesion size and shape. Previous studies have attempted to address the issue by including lesion volume as a nuisance covariate in connectome–symptom models^70, 73^. However, lesion volume captures only the overall extent of a lesion and cannot account for shape-dependent differences in which tracts are disrupted. For example, an elongated lesion and a compact spherical lesion of identical volume may intersect markedly different numbers and configurations of white-matter fibers.

The anatomy-preserving null model in rLNM offers an alternative solution. Rather than regressing out lesion volume, rLNM repositions lesions while preserving their individual size and shape, generating a size- and shape-matched null that isolates genuine tract-symptom associations from lesion-morphology effects. Using this null model, structural rLNM revealed tract-level information beyond what functional connectivity captures. For example, relative to visual deficit, visual anosognosia selectively involved a white-matter pathway linking the visual system with the hippocampus. Visual–hippocampal interactions may support recognition of visual loss by using discrepancies between expected and actual visual performance^75^. Disconnection of this pathway may therefore explain the unawareness of visual loss rather than the visual loss itself. These findings indicate that structural rLNM extends beyond functional LNM, revealing symptom-relevant tract relationships that functional connectivity alone cannot capture.

### Causal validation and translational applications of rLNM through clinical interventions

Interventional lesions from clinical surgery provide a direct causal validation paradigm for LNM. Naturally occurring lesions vary in onset, chronicity and subsequent network reorganization, such that the observed functional deficit may reflect not only the initial network disruption but also recovery and compensatory plasticity^76^. Interventional lesions, by contrast, are temporally defined and anatomically delineated, and can be paired with behavioral assessments immediately before and after intervention. In our epilepsy surgery cohort, language decline measured one week after surgery therefore provided an acute causal test of the functional consequences of disrupting the rLNM-derived language network, before long-term reorganization had occurred. The finding that greater involvement of this network by surgical resection predicted greater postoperative language decline provides patient-level interventional validation, indicating that rLNM captures a language-related network beyond lesion location per se. Clinically, rLNM-derived networks could thus serve as a network-level estimate of functional risk for surgical planning, helping to balance maximal lesion resection against preservation of critical brain function.

Beyond surgical planning, rLNM may also inform the identification of target networks for neuromodulation. Network-guided neuromodulation has increasingly used functional and structural connectivity to identify target networks for TMS^9, 77^ and DBS^74^. Selective specificity could refine the targeting paradigm by distinguishing cross-symptom and symptom-specific networks. For example, motor and visual anosognosia share impaired awareness but differ in the affected modality. Selective specificity allows neuromodulation to target the shared awareness network or the modality-specific components, depending on the intended clinical effect. This paradigm is increasingly practical as neuromodulation shifts from disorder-level toward symptom-level targeting. A recent clinical trial provides proof of principle: targeting distinct dysphoric and anxiosomatic networks differentially modulated depressive and anxious symptoms in major depression^10^. Although the candidate networks identified by rLNM require prospective stimulation-based validation, selective specificity provides a framework for precisely matching the target network to the intended therapeutic effect.

### Practical considerations for rLNM implementation

Several considerations point to opportunities for further development of rLNM. First, the anatomy-preserving null model in sensitivity testing remains computationally demanding. Although we used precomputed connectome representations and reduced the spatial resolution to 8 mm isotropic (9,473 voxels across the whole brain) to reduce computational cost (Supplementary Note 1-2, Supplementary Fig. 5-7), analyzing the amnesia dataset (43 lesions) with 10,000 permutations still required approximately 2 hours on a single workstation, with permutations executed serially. The finite number of null-model replications also limits empirical *P*-value resolution and may reduce sensitivity to small networks. Increasing this number could improve statistical power, but would entail additional computational cost. Future implementations could improve computational efficiency through parallelized computation and adaptive or accelerated permutation strategies^78^, facilitating application to substantially larger lesion cohorts and inference with more permutations. Second, the constraints in the spatial null model involve a design trade-off ^27^: highly constrained nulls may excessively restrict the space of possible randomizations, whereas an insufficiently constrained null may fail to identify genuine network convergence. The current model preserves lesion size, shape, and inter-lesion overlap. Additional functional (e.g., network degree) or biological constraints (e.g., vascular territories and etiology-specific lesion prevalence) could be incorporated as appropriate to the particular disease and hypothesis. Third, selective specificity is inherently contrast-dependent, and therefore faces the classic ‘Task B problem’ in functional neuroimaging^79^. The inferred symptom-selective components depend on the choice of comparator conditions rather than representing an absolute symptom network, so comparator selection should be hypothesis-driven and matched to the symptom-specific effects to be isolated. This places higher demands on lesion datasets, requiring carefully designed and well-characterized cohorts for selective specificity analysis. Despite these considerations, rLNM provides an open and extensible framework that enables re-evaluation of existing LNM findings and promotes greater transparency, standardization, and reproducibility across future analyses.

## Methods

### Lesion datasets

We collected lesion datasets spanning 10 neuropsychiatric symptoms from previously published studies and publicly available resources, including visual deficit^80^, amnesia^81^, stuttering^82^, migraine^83^, visual and motor anosognosia^29^, limb ataxia^84^, alien limb syndrome ^60^, addiction^71^, and neglect^80^ (see detailed information in Supplementary Table 1). The visual deficit dataset included 40 patients with visual field defects, color agnosia, or both. The amnesia dataset included 43 patients with focal brain lesions associated with clinically evident episodic memory impairment; all cases presented with anterograde amnesia and were identified through a systematic Medline search of case reports describing acquired amnesia following focal brain lesions. The stuttering dataset comprised 20 cases of acquired neurogenic stuttering following stroke. The migraine dataset included neuroimaging coordinates of decreased gray matter volume derived from a voxel-based morphometry (VBM) meta-analysis (*n* = 11 regions). The anosognosia dataset included lesions associated with visual anosognosia (*n* = 24) and motor anosognosia (*n* = 95). The limb ataxia dataset comprised 35 patients with clinically confirmed acute limb ataxia. The alien limb dataset included 48 patients with focal brain lesions associated with alien limb syndrome, identified from published case reports describing involuntary movements attributed to external control. The addiction dataset included 34 patients with focal brain lesions who were active daily smokers at the time of lesion onset. The neglect dataset included 34 patients. To provide a control condition, we constructed a mixed-lesion dataset by randomly sampling 100 lesions from the ten symptom datasets described above.

### Task activation datasets

Following the PRISMA (Preferred Reporting Items for Systematic Reviews and Meta-Analyses) guidelines (http://www.prisma-statement.org), we systematically reviewed functional neuroimaging studies from the BrainMap database (www.brainmap.org). To ensure data quality and homogeneity, we applied the following inclusion criteria across all domains: (1) use of fMRI or PET imaging; (2) involvement of healthy adult participants only (experimental context: normal mapping); (3) reporting of whole-brain results in standard stereotactic space (MNI or Talairach); and (4) inclusion of activation data only (experimental activation: activations only).

In this study, we focused on two core cognitive domains^85^: language and working memory. Language experiments were identified using ‘cognition-language’ and ‘language’ as the behavioral-domain and experimental-context criteria, respectively, yielding a final dataset of 33 papers and 155 experiments. For the working-memory analysis, we restricted the search to the well-established n-back paradigm to improve consistency across studies. To investigate shared and condition-specific effects, eligible experiments were further categorized by stimulus modality based on detailed inspection of their metadata. Only experiments involving verbal stimuli (e.g., words or sentences; *n* = 58 experiments), faces (e.g., human faces; *n* = 47 experiments) or visuospatial information (e.g., spatial locations or navigation; *n* = 79 experiments) were included in the analyses.

### Interventional lesion cohort from epilepsy surgery

To evaluate whether rLNM-derived networks capture the causal functional consequences of interventional brain lesions, we analyzed an independent multicenter cohort of 56 patients (24 females; age = 25.7 ± 9.6 years) with drug-resistant epilepsy who underwent surgical resection of epileptogenic tissue. Patients were recruited from the First Affiliated Hospital of Fujian Medical University (FAH-FMU; *n* = 37) and Sanbo Brain Hospital of Capital Medical University (SBH-CMU; *n* = 19) between 2019 and 2025. Patients with other neurological or psychiatric disorders or preoperative aphasia (WAB-AQ < 93.8, the established diagnostic cutoff^86^) were excluded. The study was approved by the institutional ethics committee at each participating center, and written informed consent was obtained from all patients or their legal representatives.

All patients underwent structural MRI and assessment with the Western Aphasia Battery Aphasia Quotient^87^ (WAB-AQ) within 1 week before surgery and again approximately 1 week after surgery. Postoperative assessments were conducted only when patients were clinically stable, showed a stable cognitive state and had no substantial residual edema on MRI or other acute postoperative complications that could confound language performance. Patients showing both a decline in WAB-AQ exceeding the reliable change index threshold^88^ (score change > 2) and a postoperative score below the established diagnostic cutoff^86^ of 93.8 were assigned to the postoperative aphasia group (*n* = 29), whereas others were assigned to the non-aphasia group (*n* = 27). For each participant, the interventional lesion was identified in the native space of the postoperative T1w images, with preoperative T1w images and postoperative T2w FLAIR images as references. Lesion boundaries were manually delineated slice by slice using MRIcron v1.0.2 software (https://www.nitrc.org/projects/mricron). All lesion masks were subsequently reviewed and confirmed by a senior neuroradiologist and a senior neurosurgeon to ensure accuracy.

### MRI data and preprocessing

Structural and resting-state fMRI data from 1,000 healthy participants in the Brain Genomics Superstruct Project (GSP)^89^ were used to construct a normative connectome. Preprocessing of both structural and functional MRI data was performed using DeepPrep^90–92^ with parameters consistent with our previous studies^93, 94^. The fMRI preprocessing pipeline included the following steps: (1) slice timing correction; (2) head motion correction; (3) linear detrending and bandpass filtering within the range of 0.01-0.08 Hz; and (4) nuisance regression, including the six head-motion parameters, white matter, ventricular and global signals, and their first-order temporal derivatives. Volumetric fMRI data were normalized to a 2-mm isotropic template (the FSL version of the MNI ICBM152 nonlinear template), followed by spatial smoothing using a 6-mm full-width at half-maximum (FWHM) isotropic Gaussian kernel within the brain mask. For analyses involving the cerebellum, the mean BOLD signal from cerebral cortical voxels within a 20-mm Euclidean radius of each cerebellar voxel was additionally regressed from that voxel’s time series before FC estimation, following established cerebellar FC procedures^95^. All lesion masks were first normalized to the same 2-mm isotropic volumetric template using a 12-degree-of-freedom linear transformation (FSL v6.0)^96^. In this study, functional rLNM analyses were conducted at an 8-mm isotropic resolution to reduce computational complexity (Supplementary Note 2, Supplementary Fig. 7). For visualization, volumetric data were resampled to 1-mm resolution and projected onto the fsaverage cortical surface using FreeSurfer^97^.

### Sensitivity testing

To control for false-positive and false-negative findings, we incorporated null model-based testing into the cLNM framework. We first constructed an LNM map from the observed lesions. Each lesion was treated as an individual region of interest (ROI), and its resting-state functional connectivity (FC) profile was estimated using normative rs-fMRI data from 1,000 healthy participants in the GSP dataset^89^. In cLNM, the mean BOLD time series within each ROI is correlated with that of every other brain voxel for each normative participant, and the resulting Fisher *r*-to-*z*-transformed FC maps are entered into voxel-wise one-sample *t*-tests. The resulting *t*-maps are FDR-corrected using the Benjamini–Hochberg procedure (*P* < 0.01), thresholded at |*t*| > 7 and binarized before calculating voxel-wise connectivity overlap across lesions. To enable computationally feasible null-model testing, we used an accelerated implementation in which lesion-wise *r*-maps were derived from the group-average GSP connectome and thresholded using a prespecified |*r*|-based criterion as in Van den Heuvel et al.^11^ (Supplementary Note 1, Supplementary Fig. 5-6). This approach has been shown to produce LNM maps highly concordant with those obtained using conventional one-sample *t*-tests^11^. After thresholding and binarization, the observed LNM map was calculated as the proportion of lesions showing suprathreshold connectivity voxel by voxel.

To assess the statistical significance of the observed LNM map, we implemented an anatomy-preserving spatial null model that constrains lesion size, shape, and spatial overlap^16^. Specifically, lesions were randomly repositioned across the brain under these constraints. A global optimization procedure based on simulated annealing^34^ was used to match the inter-lesion overlap distribution between synthetic and observed lesions, with distributional similarity assessed using a two-sample Kolmogorov-Smirnov (KS) test. For each iteration, ROI-specific *r*-maps were then computed for the synthetic lesions using the same pipeline as the observed lesions. This process was repeated 10,000 times to generate a null distribution of LNM maps. The observed LNM map was subsequently compared against this distribution using voxel-wise testing to derive a statistical *P*-map. FDR correction was applied across all 9,473 voxels in the whole-brain analysis mask (8-mm isotropic resolution; see Supplementary Note 2), and voxels with *P*_FDR_ < 0.005 were retained to define the final null model-controlled sensitivity map.

### Selective specificity testing

Selective specificity testing in rLNM aims to distinguish connectivity patterns shared across related symptoms from those selectively associated with a specific symptom. The analysis comprised two steps.

#### Cross-symptom map

A cross-symptom map was constructed to capture shared functional patterns across the symptoms included in the comparison. Specifically, lesion-wise connectivity maps were first generated using normative FC data from 1,000 participants in the GSP dataset. For each symptom, the corresponding lesion-wise maps were averaged to derive a symptom-level connectivity map. The resulting symptom-level maps were then restricted to positive connectivity and averaged voxel-wise with equal weight across symptoms. The averaged map was then masked by the intersection of the binary significance masks obtained from null-model sensitivity testing for all symptoms in the comparison, to generate the final cross-symptom map. If the intersection was empty, no significant cross-symptom region was identified.

#### Symptom-specific map

Symptom-specific effects were derived using a lesion-level regression framework followed by label-permutation testing. For each lesion, a leave-one-lesion-out (LOLO) shared-connectivity regressor was constructed by excluding the lesion, averaging the remaining lesion-wise connectivity maps within each symptom and then taking an equal-weight mean across the resulting symptom-level maps. The individual lesion connectivity map was then regressed on its corresponding LOLO shared-connectivity regressor across voxels using ordinary least squares. The resulting residual map was defined as the lesion-specific map that represented lesion-level connectivity not explained by the shared component.

For each SOI, the lesion-specific residual maps were divided into a target group comprising lesions associated with the SOI and a contrast group comprising lesions associated with all other symptoms. The two groups were compared at each voxel using Welch’s t-test. To assess significance, symptom labels were randomly reassigned across lesion-specific residual maps 10,000 times while preserving group sizes. For each permutation, the voxel-wise Welch’s *t*-statistic contrasting the SOI with the other symptoms was recalculated to generate a null distribution, against which the observed statistic was compared to obtain an empirical *P* value. Voxels with *P*_FDR_ < 0.05 were retained to define the symptom-specific map.

### Structural rLNM analysis

rLNM was extended to structural connectomes to identify symptom-relevant white-matter pathways. A normative tractogram was generated from the HCP842 atlas, a population-averaged spin distribution template reconstructed from diffusion MRI data from 842 Human Connectome Project participants^98^. Deterministic fiber tracking was performed in DSI Studio^99^ using previously described parameters^98^, yielding 550,000 whole-brain streamlines. Lesion-associated structural connectivity was then estimated using BCBToolKit^100^. For each lesion, streamlines intersecting the lesion were identified and converted into a voxel-wise streamline-count map, in which each voxel value represented the number of lesion-intersecting streamlines traversing the voxel. To retain the strongest structural connections, voxels with non-zero values were ranked by streamline count, and the top 30% were assigned a value of 1, with all remaining voxels assigned a value of 0. The resulting binary maps were aggregated across lesions of each symptom to calculate the voxel-wise proportion of lesions connected to each location.

Null lesions were generated using the same anatomy-preserving procedure as in the functional sensitivity analysis. For each of 1,000 null lesion sets, lesion-wise structural connectivity maps and their overlap were computed using the same pipeline, yielding a voxel-wise null distribution of structural convergence. Because the analysis was performed at 1-mm resolution and required repeating the complete structural-connectivity pipeline for every null set, generating the substantially larger number of permutations required for voxel-wise FDR correction was computationally prohibitive ^78^. We therefore applied a stringent uncorrected voxel-wise threshold: only voxels whose observed convergence exceeded the corresponding values in all 1,000 null lesion sets were retained to define the final structural-connectome-based sensitivity map (*P* < 10^-3^, uncorrected). To derive the selective-specificity map for an SOI, its voxel-wise sensitivity values were regressed on those of the comparison symptom, and the resulting residuals were retained to form the SOI-specific map.

### Assessment of sensitivity and specificity using simulated data

#### Simulated functional connectivity matrix

To systematically evaluate rLNM under a controlled network architecture, we simulated a whole-brain FC matrix based on a predefined modular architecture. The matrix comprised seven cortical networks defined by the Yeo atlas^101^ and a background region representing the subcortex and cerebellum. Because the simulation was designed primarily to evaluate rLNM within a cortical network architecture, interactions among subcortical nuclei and cerebellar regions were not explicitly modeled.

The matrix simulated the distribution of Fisher *z*-transformed correlation coefficients of the empirical FC matrix, and was generated according to two rules. First, within-network connections were assigned strong positive connectivity with a mean correlation of *r* = 0.30 and a standard deviation of *σ* = 0.25. All remaining connections were assigned a mean correlation of *r* = 0 and a standard deviation of *σ* = 0.15. To assess the influence of connectome hub structure on LNM results, one cortical network was designated as a global hub, and between-network correlations involving this network were increased by 0.10. Second, to emulate node-strength heterogeneity in real connectomes, we introduced continuous variation in node strength^16, 102^. Each voxel was assigned an intrinsic connectivity propensity drawn from a lognormal distribution. Each connection was then scaled by the propensities of its two voxels, such that highly connected voxels formed systematically stronger connections throughout the matrix, whereas weakly connected voxels formed weaker connections. Together with a heavy-tailed generalized normal (Laplace) per-pair noise term, this produced a continuous, long-tailed node-strength distribution.

#### Simulated lesions

To evaluate the accuracy of sensitivity and selective specificity testing in rLNM, we generated simulated lesion sets with predefined ground-truth networks. Three simulation models were used.

Lesion-proportion simulation (*α* simulation) for sensitivity testing. Five non-hub cortical networks (excluding the limbic network because of its relatively small spatial extent) were selected in turn as target networks. The lesion proportion *α* ranged from 0 to 1 in increments of 0.05, following the strategy adopted in the previous study^16^. For each value of *α*, 1000 independent lesion sets were generated, each comprising 200 lesions. A proportion *α* of the lesions was placed entirely within the target network, whereas the remaining proportion, 1 − *α*, was randomly positioned across the whole brain. Lesion size was sampled uniformly between 5 and 50 voxels.

Voxel-proportion simulation (*β* simulation) for sensitivity testing. The design followed the α simulation, except that *β* controlled the proportion of target-network voxels within each lesion.

Specifically, each lesion contained a proportion *β* of voxels from the target network and a proportion 1 − *β* of voxels from the remainder of the brain.

Shared-specific simulation (*γ* simulation) for specificity testing. The same networks used in the sensitivity simulations were included. One was designated as the cross-symptom network, and the remaining four were assigned as symptom-specific networks to four lesion groups, respectively. For each value of the shared-component proportion *γ*, each group comprised 200 lesions, with lesion sizes sampled uniformly between 10 and 50 voxels. Each lesion contained a proportion *γ* of voxels from the cross-symptom network and a proportion 1 − *γ* of voxels from its symptom-specific network. The proportion *γ* ranged from 0 to 1 in increments of 0.05.

#### Evaluation of LNM performance

Both sensitivity and specificity simulations comprised 1,000 Monte Carlo trials at each parameter level. In each trial, simulated lesion sets were generated and then analyzed using both the cLNM and rLNM pipelines. Performance was quantified using three metrics^16^. Under the complete-null condition (α/*β* = 0 or *γ* = 1), the family-wise error rate (FWER) evaluated the probability of obtaining at least one false-positive voxel, estimated as the proportion of the 1,000 trials yielding a non-empty lesion sensitivity or selective specificity map. For non-null conditions, the false-positive rate (FPR) was defined as the proportion of voxels outside the ground-truth network that were incorrectly identified as significant. The false-negative rate was defined as the proportion of voxels within the ground-truth network that were not identified as significant, and statistical power was defined as one minus the false-negative rate. The two voxel-wise metrics were summarized as the mean and standard deviation across trials.

### Prediction of language decline following surgical resection

#### Lesion anatomy-based prediction

To determine whether postoperative language decline could be explained by focal lesion anatomy, we performed voxel-based lesion-symptom mapping (VLSM)^103^ comparing the aphasia and non-aphasia groups. At each voxel, lesion involvement was compared between groups using a chi-square test. The *χ*² statistic was assigned a positive sign when lesion involvement was more frequent in the aphasia group. Multiple-comparison correction was performed using 5,000 label permutations, yielding a permutation-corrected VLSM map at *P* < 0.05. Patient-level prediction was evaluated using leave-one-subject-out cross-validation (LOSOCV). In each iteration, one patient was held out, and the uncorrected and permutation-corrected VLSM aphasia-specificity maps were recomputed using the remaining patients. The risk score of the held-out patient was calculated as the mean VLSM value within the surgical resection mask. After completing all LOSOCV iterations, predictive performance was assessed using Pearson’s correlation between the cross-validated VLSM risk scores and postoperative WAB decline. Patient-level risk scores were min-max normalized to the range 0 to 1 for comparison with the network-based measures.

#### Network-based prediction

To determine whether postoperative language decline was better captured at the network level, cLNM and rLNM were evaluated separately using the same LOSOCV framework. Within each LNM method, sensitivity maps were generated separately for the aphasia and non-aphasia groups. The aphasia sensitivity map was regressed on the non-aphasia sensitivity map across voxels using ordinary least squares, thereby removing network patterns shared between the two groups. Positive residuals were retained to generate the aphasia-specific map, with the residual values defined as voxel-wise cLNM- or rLNM-derived risk values. The risk score for the held-out subject was calculated as the mean voxel-wise risk value within the resection mask. We also evaluated prediction performance using a canonical language network derived from the Yeo atlas^101^. Each subject’s risk score was calculated as the proportion of voxels within the resection mask that overlapped the binary canonical language network mask.

### Comparative functional pattern analytics

To characterize the cognitive and behavioral associations of LNM maps, we performed a spatial meta-analytic comparison using the Neurosynth database (https://neurosynth.org/)^104^. Neurosynth provides whole-brain meta-analytic activation maps associated with 1,334 terms, which are derived from large-scale meta-analyses of neuroimaging studies. These maps characterize regional brain activation associated with cognitive and behavioral processes. In our analysis, anatomical terms (e.g., ‘insula’ and ‘cortical’) and redundant terms were excluded, restricting the analysis to cognitive and behavioral domains. Spatial similarity between LNM maps and each Neurosynth map was quantified using voxel-wise Pearson’s correlation coefficients. Statistical significance was determined after FDR correction for multiple comparisons across Neurosynth terms.

### Statistical analyses

To facilitate network-level quantification, we extended the Yeo et al. seven-network parcellation^101^ by adding the bodily-self network, a subnetwork of the AMN^39^, yielding an eight-network parcellation (see the detailed atlas in Supplementary Fig. 8a). Because this parcellation does not contain a dedicated language network, the language networks from the Gordon et al.^105^ and the Yeo et al. 17-network parcellation^101^ were used for analyses of the task-derived language sensitivity map and interventional lesions (see the detailed atlases in Supplementary Fig. 8b, c). Anatomical regions, including the inferior frontal gyrus (IFG), precuneus, anterior/posterior cingulate cortex (ACC/PCC), were defined using the Automated Anatomical Labeling 3 atlas ^106^. The superior temporal sulcus (STS) and posterior middle frontal gyrus (pMFG) were defined using the Glasser HCP-MMP1 atlas ^107^. Fusiform face area (FFA) was defined by the Activation-based Surface Atlases Project^108^. Correspondence between each surface map and cortical functional networks/anatomical regions was assessed using a rotation-based null-model permutation test (spin test) implemented in the BrainSpace toolbox^109^, with 1,000 rotations.

In the subcortex, the hippocampus was defined using the topographic atlas of Tian et al.^110^. The lateral geniculate nucleus (LGN) was defined using a probabilistic in vivo structural atlas derived from 7-T MRI^111^. In the cerebellum, the paravermis was defined by combining the anterior cerebellar lobe and vermis in the AAL3 atlas. For white matter analysis, 80 whole-brain white-matter bundles were defined using the HCP842 population-averaged tractography atlas^98^. Subcortical and white matter selectivity were assessed by comparing the mean unthresholded value of the sensitivity map within each target structure with the mean across all voxels in the remaining structures, using 1,000 spatial-autocorrelation-preserving surrogate maps generated with BrainSMASH^112^. To compare sensitivity maps, pairwise spatial similarity was quantified in volumetric space using Pearson’s correlation coefficients. Wilcoxon signed-rank tests were used to compare cLNM and rLNM separately for between-symptom similarity and similarity to the sum-of-C pattern, using matched observations.

For the epilepsy surgery cohort, age and lesion volume were compared between the aphasia and non-aphasia groups using Welch’s two-sample *t*-test. Sex distributions were compared using a χ² test of independence, with Fisher’s exact test performed as a sensitivity analysis. Differences in lesion spatial distributions between the two groups were assessed using a rotation-based null-model permutation test (spin test) implemented in the BrainSpace toolbox^109^, with 1,000 rotations. Associations between aphasia-specific risk scores and postoperative WAB decline were assessed using Pearson’s correlation, with two-sided *P* values.

## Data availability

The normative structural and resting-state functional MRI data from GSP are publicly available at https://dataverse.harvard.edu/dataverse/GSP. The normative tractogram from the HCP842 atlas is publicly available at https://brain.labsolver.org/hcp_trk_atlas.html. The cortical parcellation atlas is publicly available at https://surfer.nmr.mgh.harvard.edu/fswiki/CorticalParcellation_Yeo2011. The volumetric brain template used for visualization is an ultrahigh-resolution ex-vivo brain in MNI space^113^, which is available at https://datadryad.org/stash/downloads/file_stream/182489.

## Code availability

The code for the proposed framework is available on Github at https://github.com/pBFSLab/rLNM. Software packages incorporated into the above code for data analysis included: Matlab R2020a, https://www.mathworks.com/; Python v3.11.6, https://www.python.org; MRIcroN v1.0.2; Connectome Workbench v1.5, http://www.humanconnectome.org/software/connectome-workbench.html; Freesurfer v6.0.0, https://surfer.nmr.mgh.harvard.edu/; FSL v6.0, https://fsl.fmrib.ox.ac.uk/fsl/fslwiki.

## Acknowledgment

This work was supported by Beijing Natural Science Foundation JQ26046 (J.R.); the Changping Laboratory (H.L., J.R., W.C); the Fujian provincial recruitment project for high-level talent teams in medical and health sciences YJRCTD-2025LHS (H.L.); the Young and Middle-aged Key Personnel Training Project of Fujian Provincial Health Commission No.2024GGA024 (F.W.); and the Joint Funds for the Innovation of Science and Technology, Fujian Province No.2024Y9168 (F.W.); National Institutes of Health grants MH096773 (N.U.F.D.), MH122066 (E.M.G., N.U.F.D.), MH121276 (E.M.G., N.U.F.D.), MH124567 (E.M.G., N.U.F.D.), NS129521 (E.M.G., N.U.F.D.), and NS088590 (N.U.F.D.); the Intellectual and Developmental Disabilities Research Center (N.U.F.D.); by the Kiwanis Foundation (N.U.F.D.); and the Washington University Hope Center for Neurological Disorders (E.M.G., N.U.F.D.).

## Author contributions

Conception: H.L., J.R., W.C. Design: W.C., J.Z., Y.Long and J.R. Data acquisition, analysis and interpretation: W.C., J.Z, Y.Long, F.W., Y.Y., X.W., B.Z., Y.Lu, W.Z., X.F., J.H., L.L., G.L., Y.Lin, E.M.G., J.R. and H.L. Manuscript writing and revision: W.C., J.Z., Y.Long, F.W., E.M.G., N.U.F.D., D.W., A.Z., J.R. and H.L. All authors approved the final manuscript.

---

**Extended Data Fig. 1.**
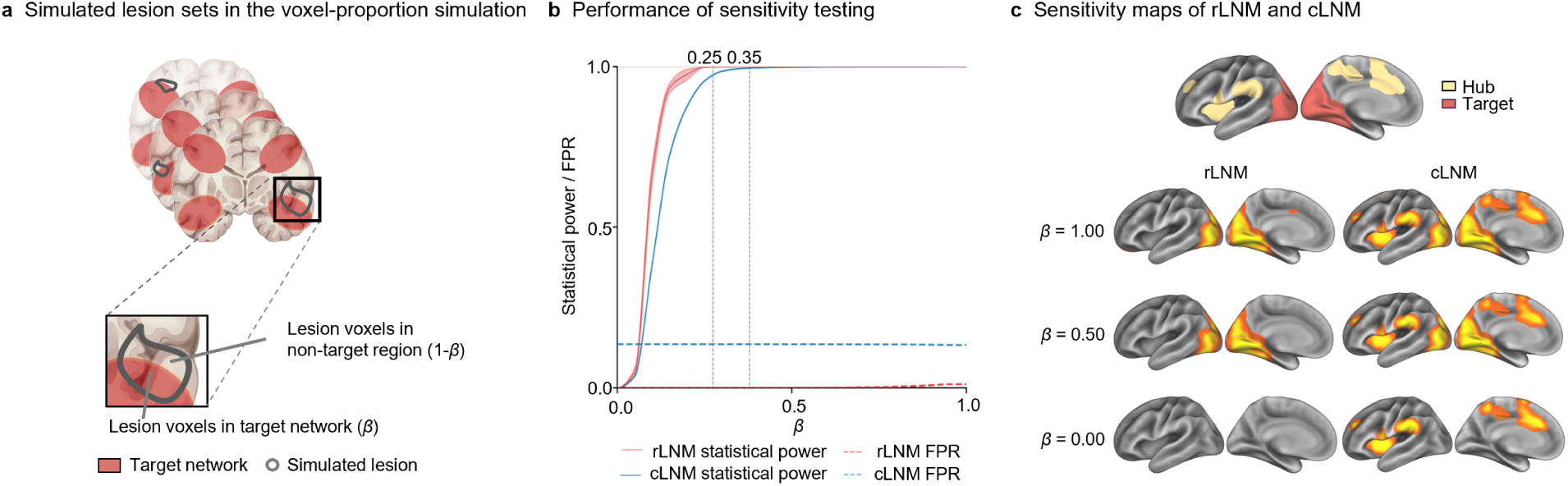
rLNM controls both false positives and negatives in voxel-proportion simulations. **a**, Generation of simulated lesion sets in the voxel-proportion simulation. For each simulated lesion (gray outline) a proportion *β* of voxels was placed within the target network (red region), and the remaining proportion, 1-*β*, was placed outside the target network. **b**, Statistical power (solid lines) and false-positive rate (FPR; dashed lines) of rLNM (red) and cLNM (blue) across *β* values ranging from 0 to 1. Statistical power reaches 1 at *β* = 0.25 for rLNM and *β* = 0.35 for cLNM. Across *β* values, the FPR of rLNM (0.0023 ± 0.0037) is consistently lower than that of cLNM (0.14 ± 0.0008; Wilcoxon signed-rank test, *P* = 4.77×10^-7^). **c**, Sensitivity maps obtained using rLNM and cLNM at representative *β* values. The top row shows the predefined hub and target networks. Subsequent rows show sensitivity maps obtained using rLNM (left) and cLNM (right) at *β* = 1.0, 0.5 and 0. Values are mean ± s.d. across simulation trials.

**Extended Data Fig. 2.**
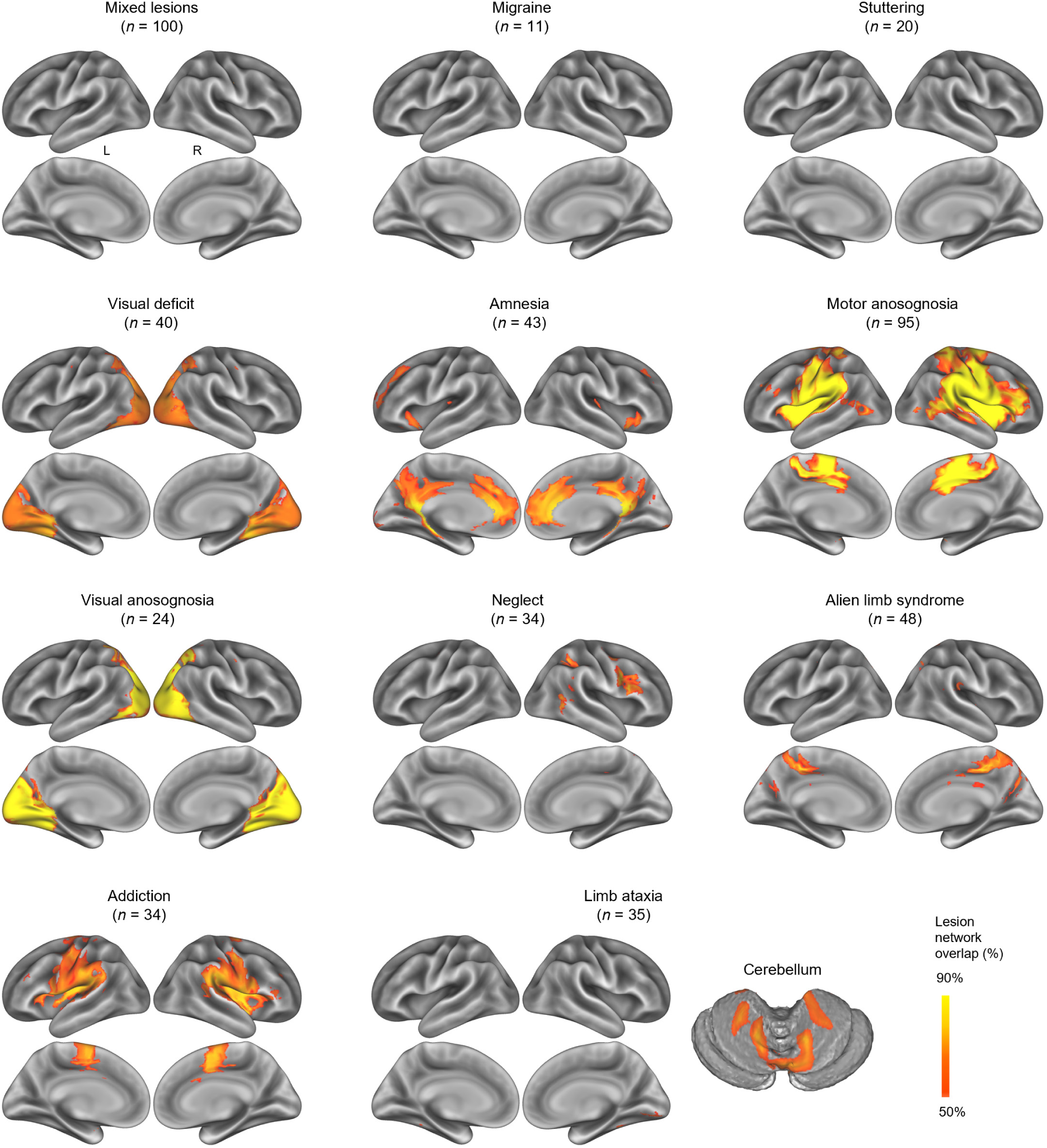
rLNM-derived sensitivity maps. Cortical surface projections show rLNM-derived sensitivity maps for mixed lesions (*n* = 100), migraine (*n* = 11), stuttering (*n* = 20), visual deficit (*n* = 40), amnesia (*n* = 43), motor anosognosia (*n* = 95), visual anosognosia (*n* = 24), neglect (*n* = 34), alien limb syndrome (*n* = 48), addiction (*n* = 34) and limb ataxia (*n* = 35). Cerebellar views are additionally shown for limb ataxia to illustrate convergence within the cerebellum.

**Extended Data Fig. 3.**
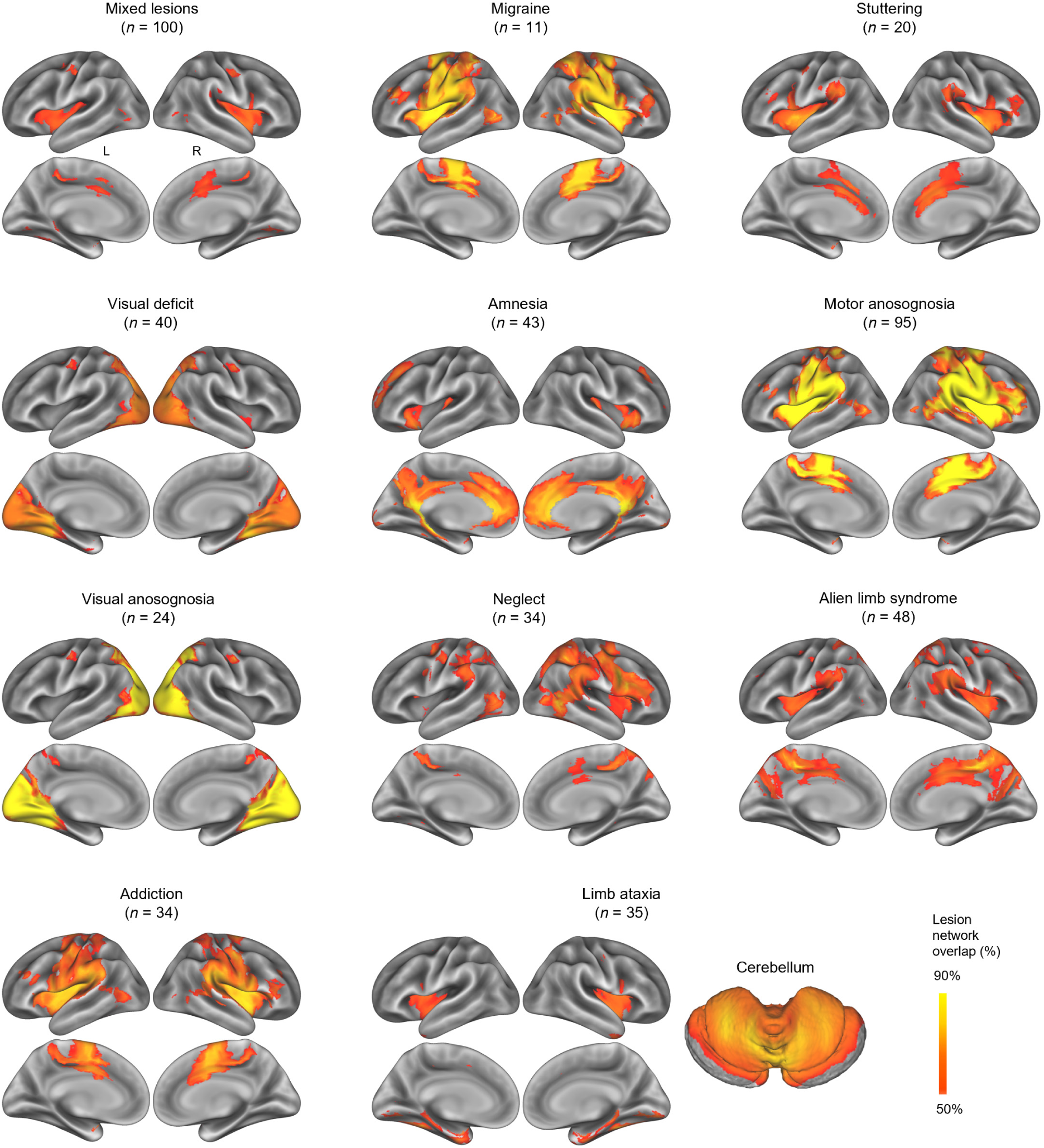
cLNM-derived sensitivity maps. Cortical surface projections show cLNM-derived sensitivity maps for mixed lesions (*n* = 100), migraine (*n* = 11), stuttering (*n* = 20), visual deficit (*n* = 40), amnesia (*n* = 43), motor anosognosia (*n* = 95), visual anosognosia (*n* = 24), neglect (*n* = 34), alien limb syndrome (*n* = 48), addiction (*n* = 34) and limb ataxia (*n* = 35). Cerebellar views are additionally shown for limb ataxia to illustrate convergence within the cerebellum. The mixed-lesion dataset was generated by randomly sampling lesions across the ten symptom datasets (*n* = 100).

**Extended Data Fig. 4.**
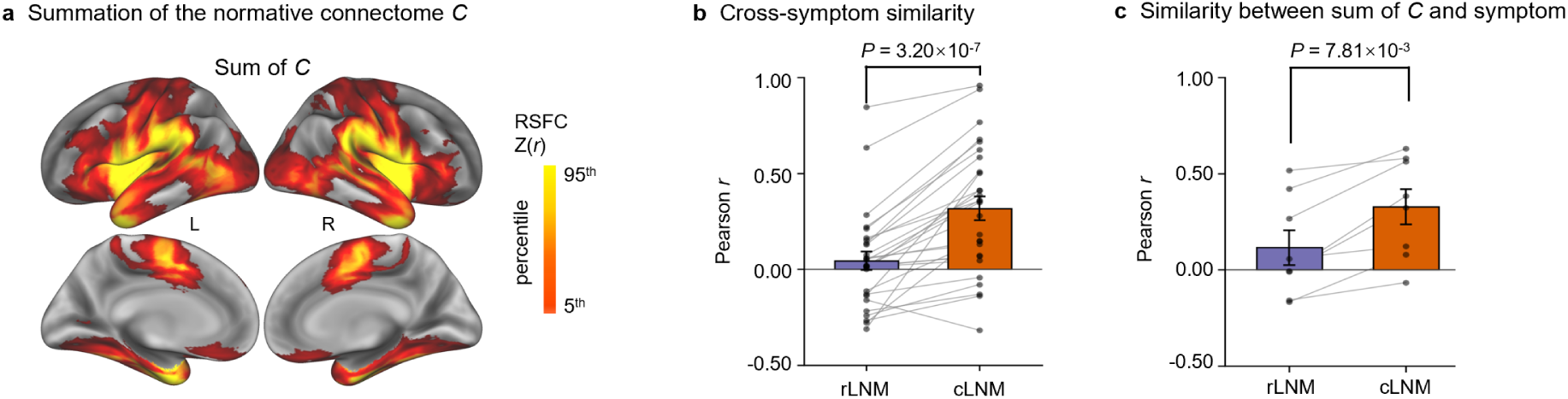
rLNM reduces between-symptom similarity and correspondence with normative connectome architecture. a, Cortical surface projection of the summed connectivity of the normative connectome (sum of *C*) derived from 1,000 healthy participants in the GSP dataset. **b,** Pairwise spatial similarity among symptom sensitivity maps generated using rLNM and cLNM. Similarity was significantly lower among rLNM maps (*r* = 0.044 ± 0.26) than among cLNM maps (*r* = 0.32 ± 0.33; Wilcoxon signed-rank test, *P* = 3.20 × 10^-7^). **c,** Spatial similarity between each symptom sensitivity map and the sum of *C*. Similarity was significantly lower for rLNM maps (*r* = 0.12 ± 0.26) than for cLNM maps (*r* = 0.33 ± 0.26; Wilcoxon signed-rank test, *P* = 7.81 × 10^-3^). Migraine and stuttering were excluded from these comparisons because no voxels were identified in their rLNM maps.

**Extended Data Fig. 5.**
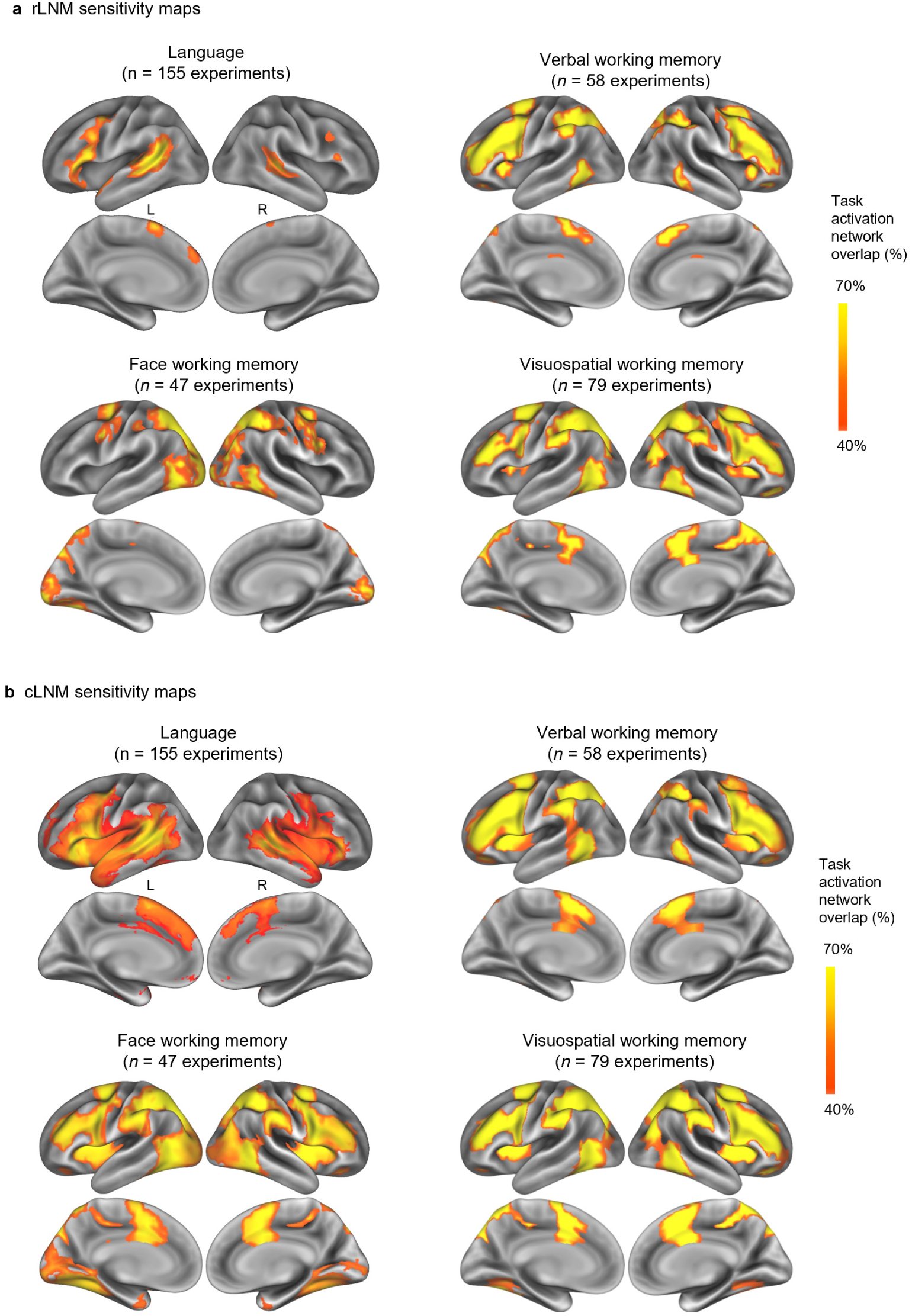
Comparison of rLNM- and cLNM-derived sensitivity maps across tasks. a, Cortical surface projections of rLNM-derived sensitivity maps for language (*n* = 155 experiments), verbal working memory (*n* = 58 experiments), face working memory (*n* = 47 experiments) and visuospatial working memory (*n* = 79 experiments). **b,** Corresponding cLNM-derived sensitivity maps for the same four task conditions.

**Extended Data Fig. 6.**
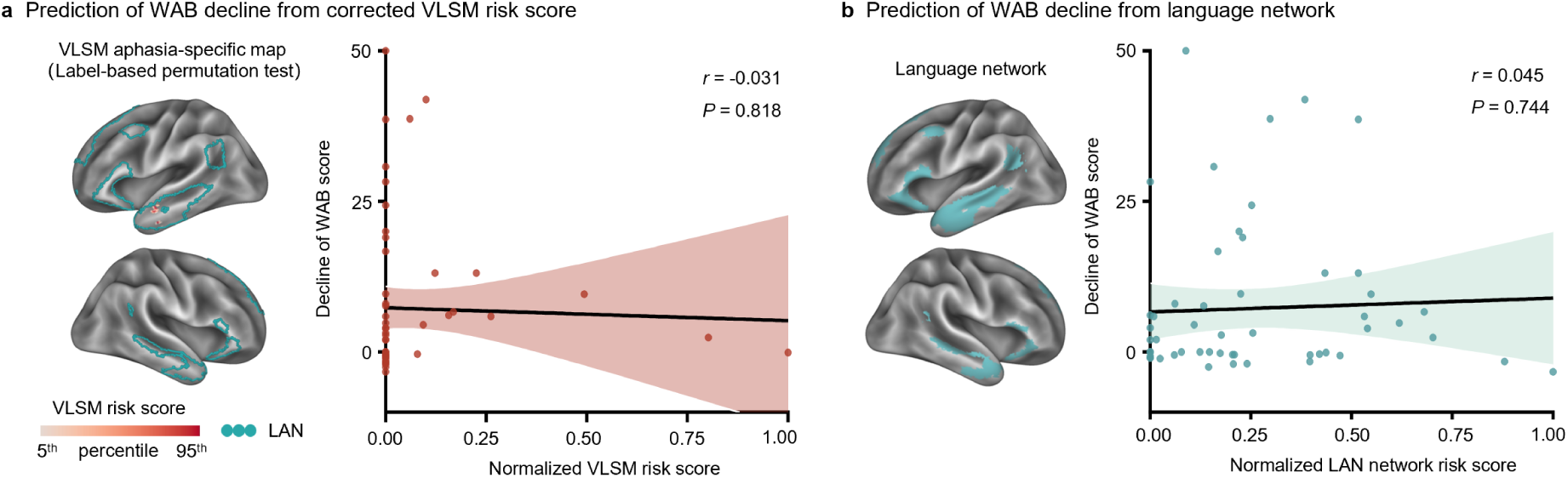
Permutation-corrected VLSM and canonical language-network involvement did not predict postoperative language decline. a, Aphasia-specific map generated using VLSM with label-permutation correction. The map was obtained by contrasting voxel-wise lesion involvement between the aphasia and non-aphasia groups. Under leave-one-subject-out cross-validation, risk scores derived from this map were not significantly associated with postoperative Western Aphasia Battery (WAB) decline (*r* = -0.031, CI = [-0.29, 0.23], *P* = 0.818). The color scale spans the 5^th^–95^th^ percentiles of the map values; cyan contours delineate the canonical language network (LAN). **b,** Canonical LAN and its association with postoperative WAB change. LAN risk scores were not significantly associated with postoperative change (*r* = 0.045, CI = [-0.22, 0.30], *P* = 0.744). Shaded regions denote 95% confidence intervals around the fitted regression lines.

**Extended Data Fig. 7.**
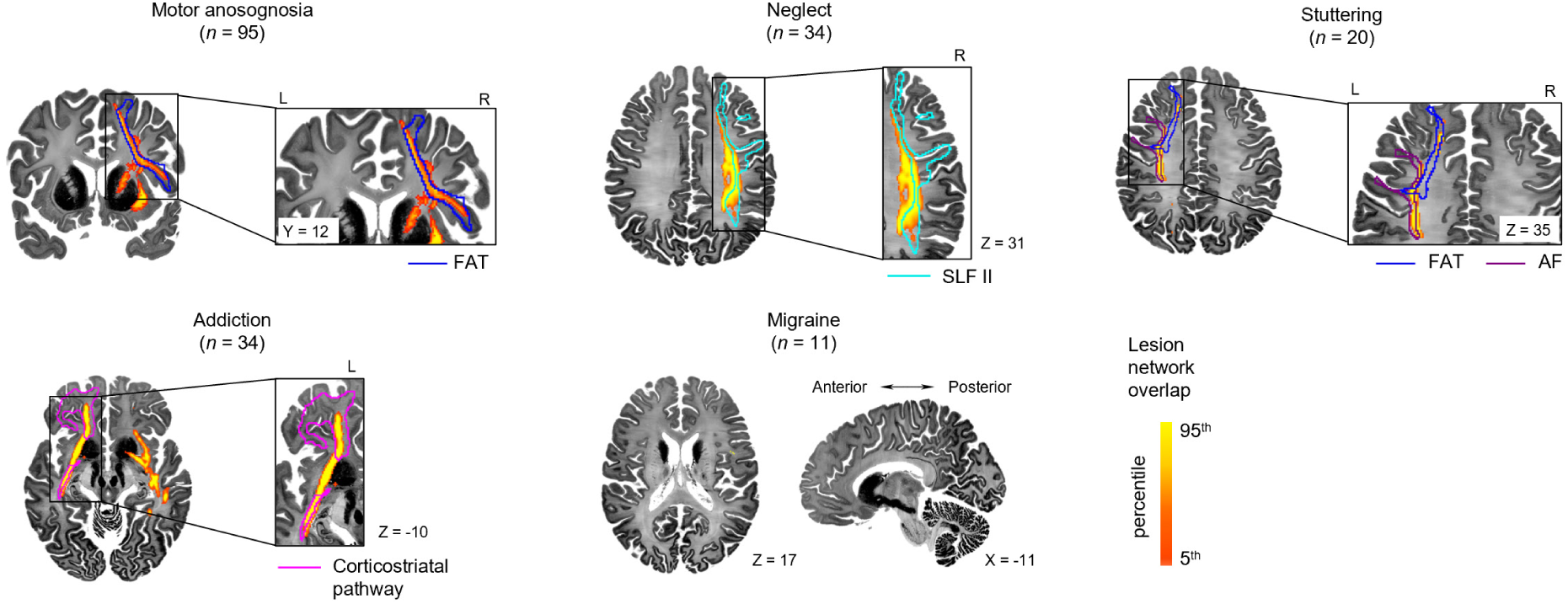
Structural rLNM sensitivity maps across neuropsychiatric symptoms. Structural rLNM sensitivity maps are shown for motor anosognosia (*n* = 95), neglect (*n* = 34), stuttering (*n* = 20), addiction (*n* = 34) and migraine (*n* = 11). Warm-colored overlays indicate lesion network overlap, expressed as percentile ranks from the 5^th^ to the 95^th^ percentile. Enlarged views show regions of high rLNM sensitivity in relation to representative white-matter tracts. The motor anosognosia map predominantly overlapped the right frontal aslant tract (FAT; one-sided spatial permutation test, 1,000 permutations, *P*_perm_ = 0.008; blue), the neglect map overlapped the right superior longitudinal fasciculus II (SLF II; *P*_perm_ = 0.003; cyan), and the stuttering map overlapped both the left FAT (blue) and arcuate fasciculus (AF; *P*_perm_ = 0.003; purple). The addiction map overlapped the corticostriatal pathway (*P*_perm_ < 10^-3^; magenta).

**Extended Data Fig. 8.**
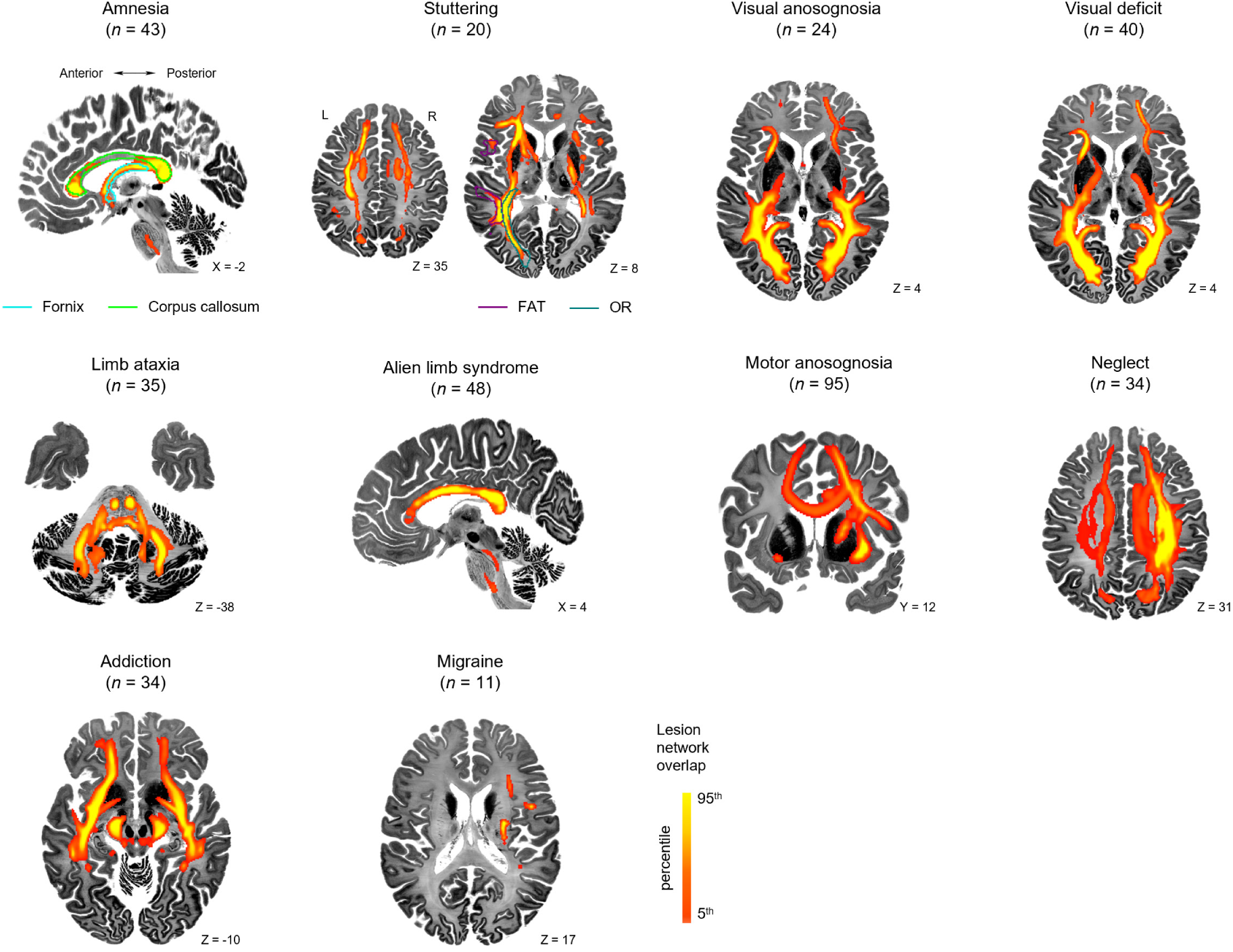
Structural cLNM sensitivity maps across neuropsychiatric symptoms. Structural cLNM sensitivity maps are shown for amnesia (*n* = 43), stuttering (*n* = 20), limb ataxia (*n* = 35), alien limb syndrome (*n* = 48), motor anosognosia (*n* = 95), neglect (*n* = 34), visual anosognosia (*n* = 24), visual deficit (*n* = 40), addiction (*n* = 34) and migraine (*n* = 11). For anatomical reference, the fornix (cyan) and corpus callosum (lime) are overlaid on the amnesia map, whereas the frontal aslant tract (FAT; magenta) and optic radiation (OR; green) are overlaid on the stuttering map. Warm-colored overlays indicate structural lesion-network convergence, displayed as percentile ranks from the 5^th^ to the 95^th^ percentile.

