## Supplementary Materials for "Robust lesion network mapping reveals symptom-specific circuits"

### Online Supplementary Materials

**Supplementary Note 1 | Approximating t-statistic thresholding using percentile-based correlation thresholding.**

**Supplementary Note 2 | Spatial downsampling for computational efficiency.**

**Supplementary Fig. 1 | Simulated functional connectivity matrix.**

**Supplementary Fig. 2 | Effects of overlap threshold on cLNM performance.**

**Supplementary Fig. 3 | Functional-network correspondence of cross-symptom and symptom-specific anosognosia maps.**

**Supplementary Fig. 4 | Demographic and lesion characteristics of the postoperative aphasia and non-aphasia groups.**

**Supplementary Fig. 5 | Consistency between group-mean *r* and group-level *t* connectivity estimates.**

**Supplementary Fig. 6 | Robustness of lesion network maps across functional-connectivity thresholds.**

**Supplementary Fig. 7 | Consistency of lesion network maps across spatial resolutions.**

**Supplementary Fig. 8 | Cortical functional network parcellations.**

**Supplementary Table 1 | Summary of lesion datasets across 10 neuropsychiatric symptoms.**

**Supplementary Table 2 | The number of surviving voxels with positive FC in rLNM sensitivity maps at MNI-8mm resolution.**

**Supplementary Table 3 | Correspondence between symptom-specific white-matter pathways and previous neuroanatomical evidence.**

**Supplementary Note 1. Approximating *t*-statistic thresholding using percentile-based correlation thresholding**

In conventional lesion network mapping (cLNM), each lesion is used as a seed to generate a functional connectivity (FC) map for each participant in a normative cohort (e.g., 1,000 subjects from the GSP dataset). For each lesion–voxel pair, FC values are tested across participants using a one-sample t-test. The resulting t-map is thresholded (commonly $|t| \geq 5$ or 7) and binarized, and the binary maps are aggregated across lesions to quantify voxel-wise connectivity overlap. To reduce computational cost, we used an accelerated implementation based on the group-average FC matrix, similar to that used by Van den Heuvel et al.[1]. For each lesion, its FC map was obtained directly by averaging the group-level connectivity profiles of its constituent voxels. The strongest positive connections were then retained using a prespecified percentile threshold and binarized to indicate whether the lesion was functionally connected to each voxel. This implementation preserves the binarization step while replacing cohort-level *t*-statistic thresholding with direct thresholding of lesion-seeded connectivity derived from the group-average connectome. This differs from approaches that operate directly on unthresholded continuous connectivity values[1].

To validate this approximation, we generated a group-average *r* matrix and a one-sample *t* matrix from the same GSP cohort. The two matrices showed strong element-wise correspondence (Pearson $r = 0.972$; Supplementary Fig. 5a). We then compared the resulting binary matrices across candidate thresholds. Retaining the top 25% of positive correlations in the *r* matrix yielded a pattern closely matching that obtained at $t \geq 7$ (Dice coefficient = 0.952; Supplementary Fig. 5b). Thus, within the GSP cohort, percentile-based thresholding provided a close approximation to conventional *t*-statistic thresholding. By avoiding repeated participant-level voxel-wise testing, this implementation substantially reduced computational demands, particularly for null-model analyses. In addition, we further assessed robustness to the choice of percentile-based *r* threshold by retaining the strongest 15–30% of positive correlations. Lesion network maps remained highly consistent across thresholds for both amnesia (mean pairwise $r = 0.96 \pm 0.02$) and visual deficits ($r = 0.98 \pm 0.01$; Supplementary Fig. 6).

**Supplementary Note 2. Spatial downsampling for computational efficiency**

In fMRI analyses, FC is commonly computed in a standard 2 mm isotropic space. In the present study, we performed analyses at a lower spatial resolution (8 mm) to reduce computational complexity. To evaluate the effect of this downsampling, we compared lesion network maps generated at 8-mm and 2-mm resolutions for two representative datasets (Supplementary Fig. 7). The maps showed strong spatial correspondence for both amnesia ($r = 0.809$, $P_{spin} < 10^{-3}$) and visual deficit ($r = 0.832$, $P_{spin} < 10^{-3}$), indicating that their large-scale spatial patterns were preserved after downsampling.

The 8-mm representation substantially decreases the number of voxels and associated connectivity calculations, yielding an approximately 400-fold speedup relative to 2-mm resolution and making extensive null-model iterations computationally feasible. This gain in efficiency comes at the cost of reduced sensitivity to fine-grained anatomical organization, particularly within small subcortical nuclei. Future studies aimed at resolving finer subcortical anatomy may therefore benefit from higher-resolution, region-specific or multiscale analyses.

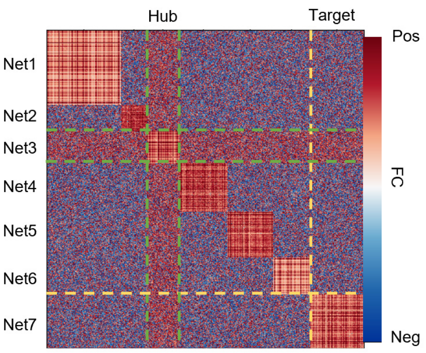


**Supplementary Fig. 1 | Simulated functional connectivity matrix.** Simulated functional connectivity (FC) matrix comprising seven cortical networks, with within- and between-network connectivity distributions informed by an empirical FC matrix. Within-network connectivity was sampled with a mean of 0.30 (s.d. = 0.25), whereas between-network connectivity had a mean of 0 (s.d. = 0.15). Connections between the designated hub network and all other networks were increased by 0.10. Dashed lines delineate the designated hub network (green) and an example target network (yellow). Blue and red indicate lower and higher connectivity, respectively.

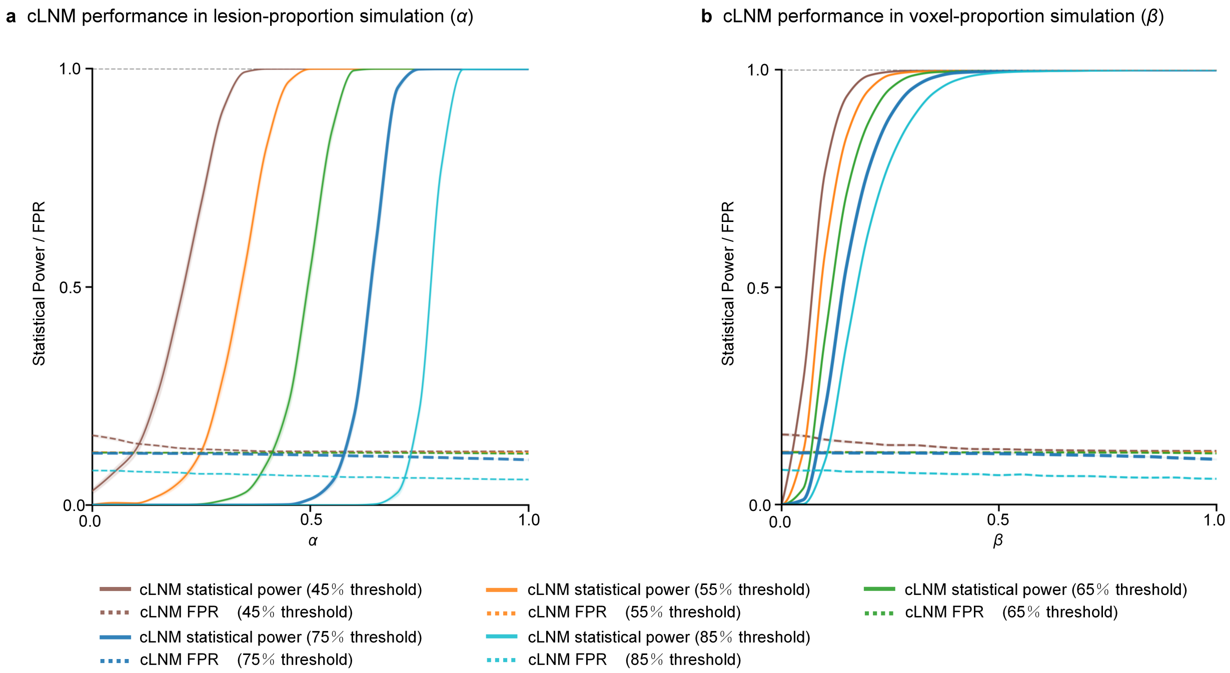


**Supplementary Fig. 2 | Effects of overlap threshold on cLNM performance.** Statistical power (solid lines) and false-positive rate (FPR; dotted lines) across fixed overlap thresholds ranging from 45% to 85% in **a**, lesion-proportion simulations across $\alpha$, and **b**, voxel-proportion simulations across $\beta$.

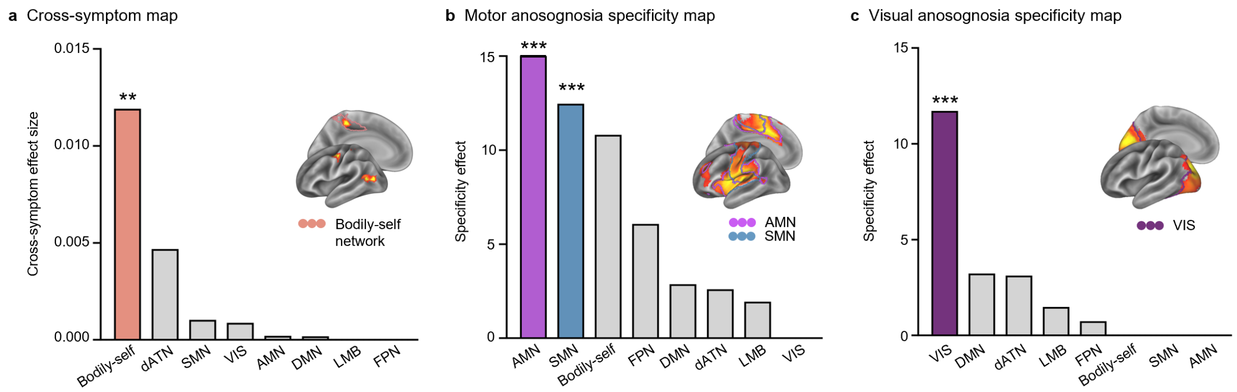


**Supplementary Fig. 3 | Functional-network correspondence of cross-symptom and symptom-specific anosognosia maps.** **a**, The cross-symptom map of motor ($n$ = 95) and visual ($n$ = 24) anosognosia corresponded significantly to the bodily-self network (spin test, 1,000 permutations, one-sided $P_{spin}$ = 0.007; all other networks, $P_{spin} > 0.05$). **b,** The specificity map of motor anosognosia corresponded to the action-mode network (AMN) and somatomotor network (SMN, all $P_{spin} < 0.001$; all other networks, $P_{spin} > 0.05$). **c**, The specificity map of visual anosognosia corresponded selectively to the visual network (VIS, $P_{spin} < 0.001$; all other networks, $P_{spin} > 0.05$). ***$P_{spin} < 0.001$, **$P_{spin} < 0.01$. dATN, dorsal attention network; FPN, frontoparietal network; DMN, default mode network; LMB, limbic network.

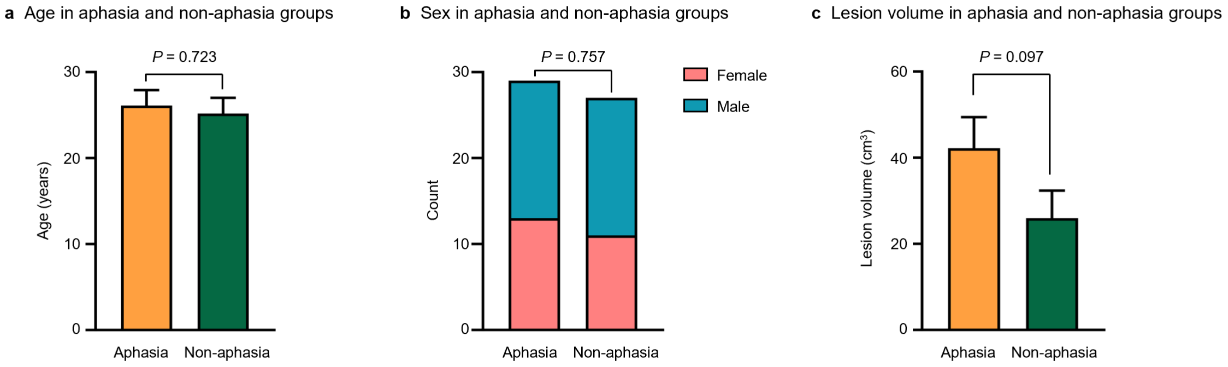


**Supplementary Fig. 4 | Demographic and lesion characteristics of the postoperative aphasia and non-aphasia groups. a,** Age did not differ between the aphasia ($n$ = 29; 26.10 ± 9.78 years) and non-aphasia ($n$ = 27; 25.19 ± 9.53 years) groups (two-sided Welch's $t$-test, $P$ = 0.723). **b,** Sex distribution was comparable between the aphasia (13 females and 16 males) and non-aphasia (11 females and 16 males) groups (two-sided Pearson's $\chi^2$ test, $P$ = 0.757; Fisher's exact test, $P$ = 0.793). **c,** Lesion volume did not differ significantly between the aphasia (42.24 ± 38.68 $cm^3$) and non-aphasia (25.97 ± 33.32 $cm^3$) groups (two-sided Welch's $t$-test, $P$ = 0.097). Values in **a** and **c** are reported as mean ± s.d.; error bars indicate s.e.m.

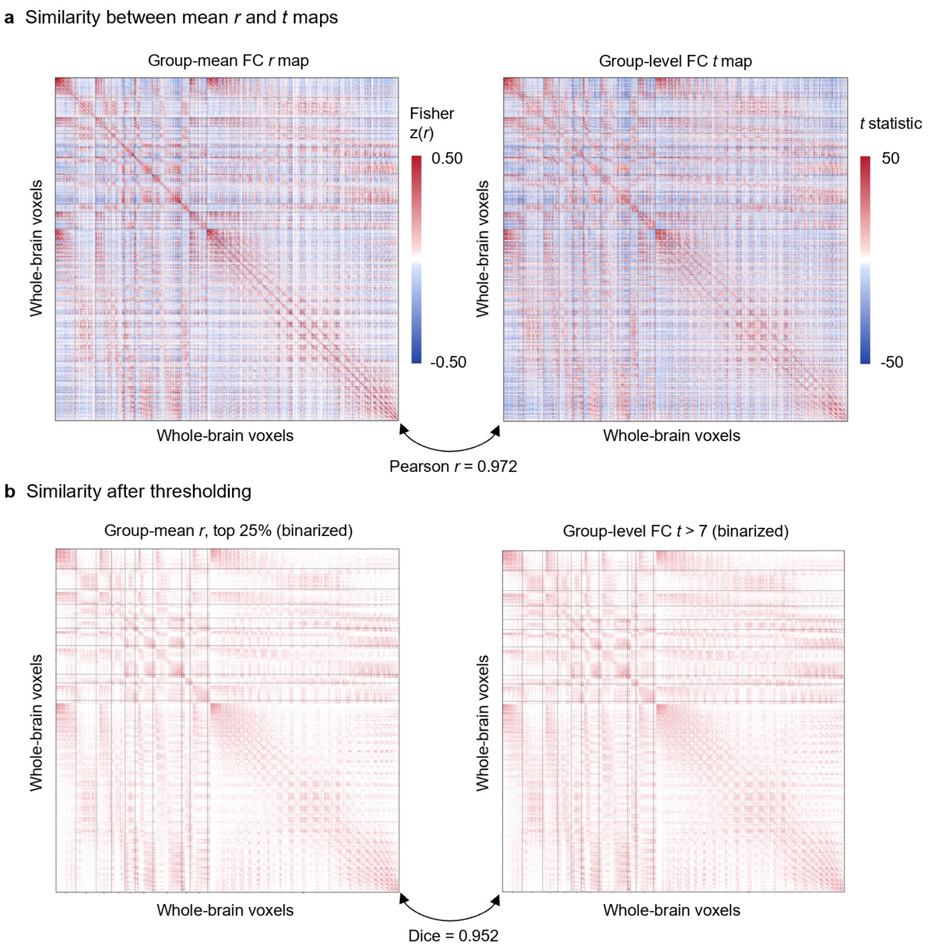


**Supplementary Fig. 5 | Consistency between group-mean *r* and group-level *t* connectivity estimates. a**, Whole-brain voxel-wise functional connectivity matrices represented as the mean Fisher *z*-transformed connectivity (*r*; left) and group-level *t* statistic (right) based on normative connectomes derived from 1,000 healthy participants in the Brain Genomics Superstruct Project (GSP) dataset[2]. Spatial correspondence between the two unthresholded matrices was quantified using Pearson correlation ($r = 0.972$). **b**, Thresholded and binarized connectivity matrices obtained by retaining the top 25% of group-mean *r* values (left) and connections with group-level $t > 7$ (right). Overlap between the resulting binary connectivity patterns was quantified using the Dice coefficient (Dice = 0.952).

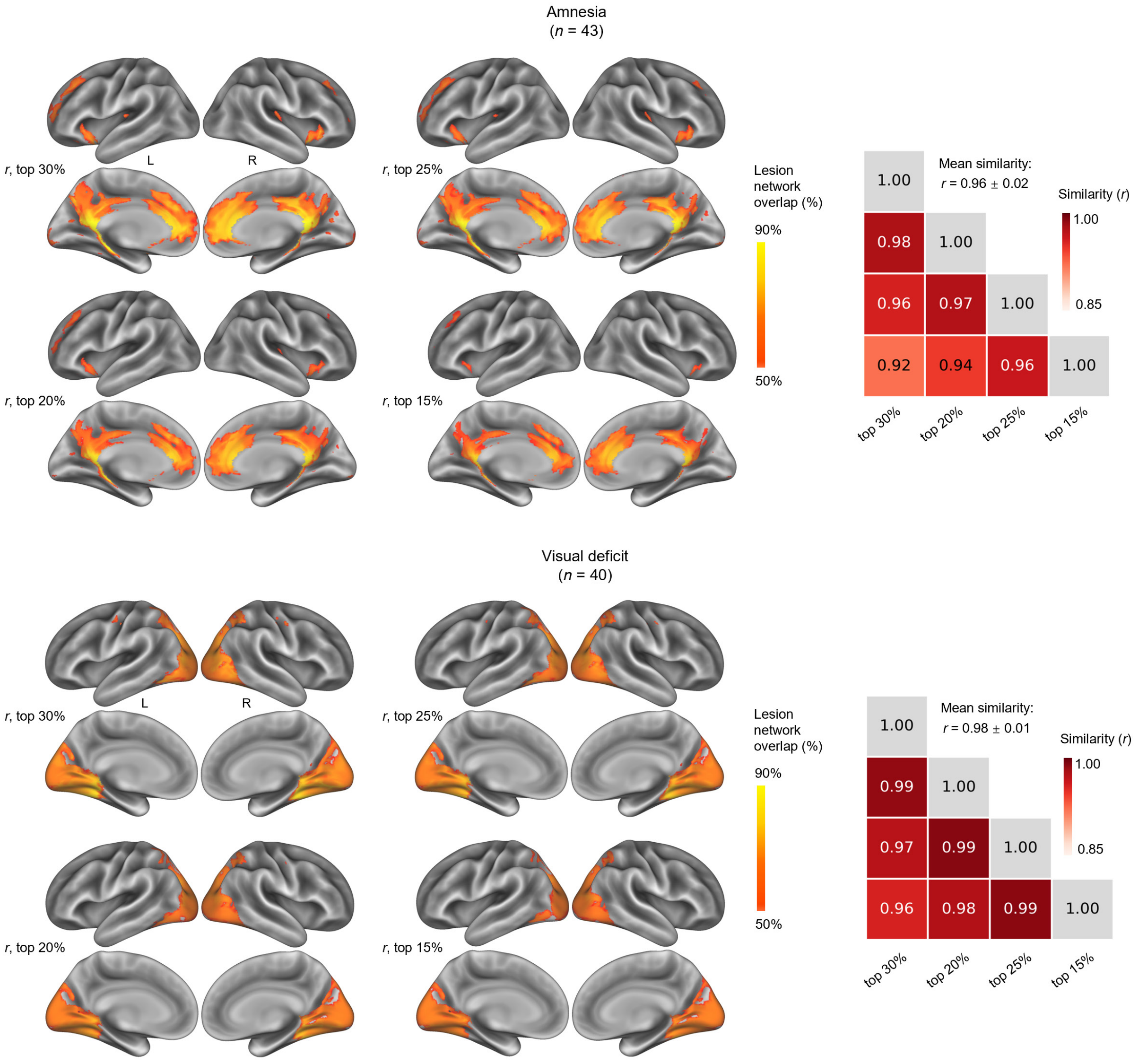


**Supplementary Fig. 6 | Robustness of lesion network maps across functional-connectivity thresholds.** Lesion network maps for amnesia (*n* = 43; top) and visual deficits (*n* = 40; bottom) were generated using percentile-based thresholds applied to each lesion-wise correlation (*r*) map. The strongest 30%, 25%, 20% or 15% of positive *r* values were retained and binarized before calculating cross-lesion network overlap. Matrices on the right show pairwise spatial similarity between maps obtained at different thresholds, quantified using Pearson correlation. Mean pairwise similarity was $r = 0.96 \pm 0.02$ for amnesia and $r = 0.98 \pm 0.01$ for visual deficit. Values are mean ± s.d.

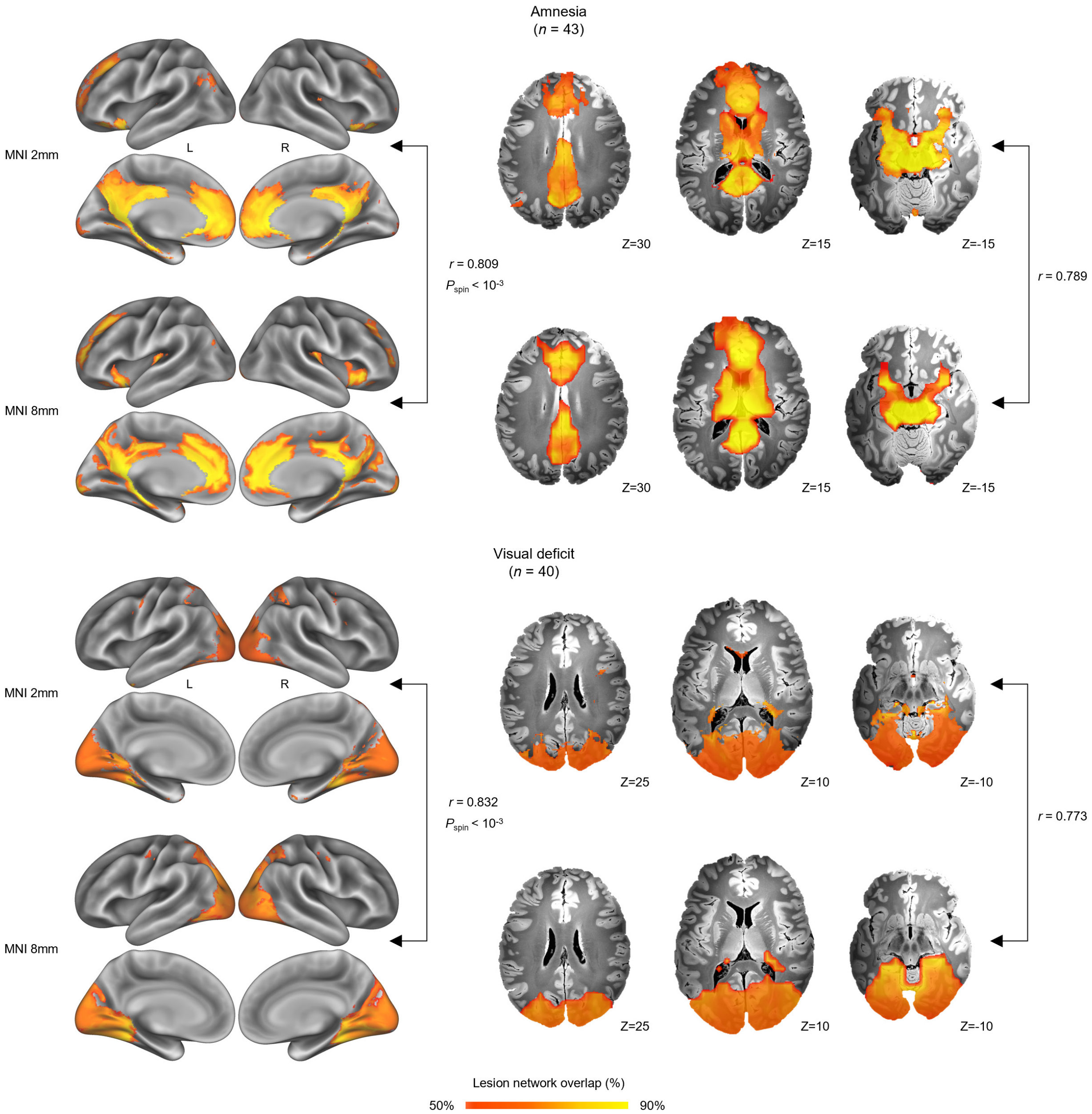


**Supplementary Fig. 7 | Consistency of lesion network maps across spatial resolutions.** Lesion network maps for amnesia ($n = 43$; top) and visual deficit ($n = 40$; bottom) generated in MNI 2-mm and MNI 8-mm spaces. Cortical surface (left) and axial views (right) show lesion network overlap. Maps derived at the two resolutions showed high spatial similarity on cortical surfaces (amnesia, $r = 0.809$; spin test, 1,000 permutations, one-sided $P_{\text{spin}} < 10^{-3}$; visual deficit, $r = 0.832$, $P_{\text{spin}} < 10^{-3}$) and in volumetric space (amnesia, $r = 0.789$; visual deficit, $r = 0.773$).

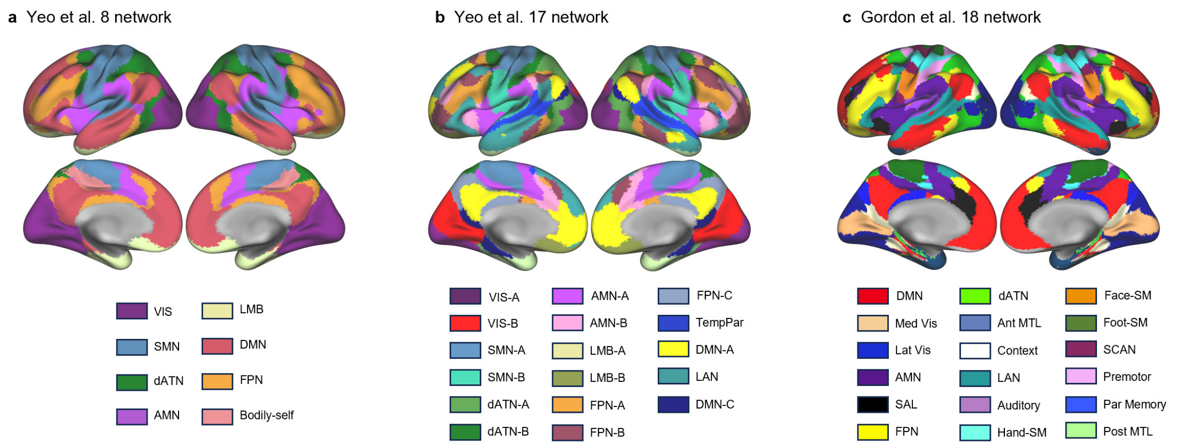


**Supplementary Fig. 8 | Cortical functional network parcellations. a**, Modified Yeo seven-network cortical parcellation. The original resting-state functional connectivity (RSFC)-derived seven-network atlas of the cerebral cortex[3] was extended by adding the bodily-self network, a subnetwork of the action-mode network (AMN)[4], yielding an eight-network parcellation. **b,** RSFC-derived seventeen-network cortical parcellation from Yeo et al.[3]. **c,** RSFC-derived eighteen-network cortical parcellation from Gordon et al.[5].

**Supplementary Table 1 | Summary of lesion datasets across 10 neuropsychiatric symptoms.**

| Dataset | No. of lesions | Symptom / phenotype | Data source and lesion definition | Primary Reference |
|---|---|---|---|---|
| Visual deficit | 40 | Visual deficits | Patients with focal brain lesions associated with visual field defects, color agnosia, or both; cases obtained from published lesion reports compiled through LesionBank. | Turner, J., et al. [6] |
| Amnesia | 43 | Episodic memory impairment / amnesia | Focal brain lesions associated with clinically evident episodic memory impairment; cases identified from published reports of acquired amnesia. | Ferguson, M.A., et al. [7] |
| Stuttering | 20 | Acquired neurogenic stuttering | Focal stroke lesions associated with acquired neurogenic stuttering. | Theys, C., et al. [8] |
| Migraine | 11 | Migraine | Meta-analytic coordinates of decreased gray-matter volume associated with migraine, derived from voxel-based morphometry studies. | Burke, M.J., et al. [9] |
| Visual anosognosia | 24 | Unawareness of visual deficit | Focal lesions associated with visual loss without awareness of the visual deficit (visual anosognosia/Anton syndrome). | Kletenik, I., et al. [10] |
| Motor anosognosia | 95 | Unawareness of motor deficit | Focal lesions associated with weakness without awareness of the motor deficit. | Kletenik, I., et al. [10] |
| Limb ataxia | 35 | Acute limb ataxia | Stroke lesions in patients with clinically confirmed new-onset acute limb ataxia. | Liesmäki, O., et al. [11] |
| Alien limb syndrome | 48 | Alien limb syndrome | Focal brain lesions associated with involuntary limb movements accompanied by impaired sense of agency or external attribution. | Darby, R.R., et al. [12] |
| Addiction | 34 | Addiction remission after brain lesion | Focal brain lesions in active daily smokers associated with abrupt remission of smoking addiction. | Joutsa, J., et al. [13] |
| Neglect | 34 | Spatial neglect | Focal brain lesions associated with neglect, obtained from published lesion cases compiled through LesionBank. | Turner, J., et al. [6] |

**Supplementary Table 2 | The number of voxels surviving FDR-corrected null model tests in rLNM sensitivity maps at MNI-8mm resolution.**

| Dataset | # surviving voxels, *n* |
|---|---|
| Visual deficit | 1964 |
| Amnesia | 1091 |
| Stuttering | 0 |
| Migraine | 0 |
| Visual anosognosia | 1984 |
| Motor anosognosia | 2330 |
| Limb ataxia | 202 |
| Alien limb syndrome | 349 |
| Addiction | 1056 |
| Neglect | 259 |

**Supplementary Table 3 | Correspondence between symptom-specific white-matter pathways and previous neuroanatomical evidence.**

| Dataset | White-matter pathway identified in the study | Anatomical and functional relevance | Primary References |
|---|---|---|---|
| Motor anosognosia | Right frontal aslant tract (FAT) | The FAT connects the inferior frontal cortex with the pre-supplementary and supplementary motor areas. The right FAT is part of a premotor disconnection network implicated in anosognosia for hemiplegia. | Pacella, V., et al. [14] |
| Neglect | Right superior longitudinal fasciculus II (SLF II) | The SLF II connects posterior parietal and frontal regions involved in spatial attention. Damage to the right SLF II has been strongly associated with spatial neglect | Thiebaut de Schotten, M., et al. [15] |
| Stuttering | Left frontal aslant tract (FAT) and arcuate fasciculus (AF) | The left FAT contributes to speech initiation, sequencing, and fluent speech production. The AF connects posterior temporal language regions with frontal speech-motor regions and supports phonological processing and auditory-motor integration. | Neef, N.E., et al. [16] Kemerdere, R., et al. [17]; Sommer, M., et al. [18] |
| Addiction | Corticostriatal pathway | Corticostriatal pathways integrate affective and cognitive-control signals involved in reward-guided action selection. Altered frontostriatal white-matter integrity has been reported in substance dependence. | Larsen, B., et al. [19]; Ghosh, A., et al. [20] |
